# Evolutionary patterns of the chimerical retrogenes in Oryza

**DOI:** 10.1101/573501

**Authors:** Yanli Zhou, Huazhi Song, Jinghua Xiao, Qifa Zhang, Manyuan Long, Chengjun Zhang

## Abstract

Chimerical retroposition delineate a process by which RNA reverse transcribed integration into genome accompanied with recruiting flanking sequence, which is asserted to play essential roles and drive genome evolution. Although chimerical retrogenes hold high origination rate in plant genome, the evolutionary pattern of retrogenes and their parental genes are not well understood in rice genome. In this study, using maximum likelihood method, we evaluated the substitution ratio along lineages of 24 retrogenes and parental gene pairs to retrospect the evolutionary patterns. The results indicate that some specific lineages in 7 pairs underwent positive selection. Besides the rapid evolution in the initial stage of new chimerical retrogene evolution, an unexpected pattern was revealed: soon or some uncertain period after the origination of new chimerical retrogenes, their parental genes evolved rapidly under positive selection, rather than the rapid evolution of the new chimerical retrogenes themselves. This result lend support to the hypothesis that the new copy assistant the function evolution among parental gene and retrogene. Transcriptionally, we also found that one retrogene (RCG3) have a high expression at the period of calli infection which supported by chip data while its parental gene doesn’t have. Finally, by calibration to Ka/Ks analysis results in other species including *Apis mellifera*, we concluded that chimerical retrogenes are higher proportionally positive selected than the regular genes in the rice genome.

## Introduction

Retroposed duplicate genes, retrogenes, result from the process of retrotransposition, in which mRNAs are reverse-transcribed into cDNA and then inserted into a new genomic position (Zhang, Wu, et al. 2005). Because of the processed nature of mRNAs, the newly duplicated paralogs lack introns, have a poly-A tail and short flanking repeats, causing function inefficiency of retrogenes for the lack of regulation element. However, chimerical retrogenes resurrect gene integrity by recruiting genome resided flanking sequence, is tenable to confer new functions and thus contribute to adaptive evolution.

The gene *Jingwei*, which originated by the insertion of a retrocopy of the Alcohol dehydrogenase gene (*Adh*) into the *yande* in *Drosophila*, was the first characterized young chimerical gene (Long and Langley 1993). Since then, many new retrogenes with chimerical structures have been reported in animals. The *Sdic* gene fused from *Cdic* and *AnnX* (Nurminsky, et al. 1998), non-protein-coding RNA gene *sphinx* (Wang, et al. 2002), retroposed fission gene family monkey king (Wang, et al. 2004) and *siren* gene derived from *Adh* (Nozawa, et al. 2005). Recently, 14 chimerical genes were identified in *Drosophila* (Rogers, et al. 2009) and one of them named *Qtzl* was observed to have male-reproductive function (Rogers, et al. 2010). It was also reported that approximately twenty retrogenes in primates and mammals (Kaessmann, et al. 2009). For example, TRIM5-CypA fusion protein (*TRIMCyp*) gene is formed by a cyclophilin A (CypA) cDNA transposed into the *TRIM5* locus (Virgen, et al. 2008; Wilson, et al. 2008); Marques worked out that approximately 57 retrogenes in the human genome emerged in primates (Marques, et al. 2005). Despite these plentiful findings of retrogenes in animals, however, no retrogenes have been systematically identified in plant until the retroposons excavation in *Arabidopsis* (Zhang, Wu, et al. 2005). Soon later, chimerical retrogenes were creative mentioned in rice (Wang, et al. 2006). In rice genome, the abundant retroposition mediated chromosomal rearrangements resulted in 898 presumed retrogenes, 380 of which were found to create chimerical gene structures, by recruiting nearby exon-intron sequences. Many of these chimerical retrogenes originated recently, while how did they shape their fortunes are poorly understood.

Since the searching of new retrogenes becomes technically easier, more opportunities are available to further investigate the evolutionary patterns of chimerical retrogenes. Parallel changes in the spatial and physicochemical properties of functionally important protein regions, have been reported in the evolution of young chimerical genes (Zhang, et al. 2004). Three retrogenes in *Drosophlia*, i.e., *Jinwei, Adh-Finnegan* and *Adh-Twain*, were found to undergo rapid adaptive amino acid evolution in a short period of time after they were formed, then followed by later quiescence and functional constraint (Jones and Begun 2005; Jones, et al. 2005). The finding of the initially-elevated and subsequent slowdown substitution pattern concluded the first insight into the adaptive evolutionary process of the new genes.

Although the rice genomes have a high rate to generate chimerical retrogenes, the patterns of sequence evolution and underlying mechanisms to prompt these new retrogenes are unclear. To understand these two critical aspects of new gene evolution, we analyzed 24 retrogenes by choosing randomly from 380 chimerical retrogenes suggested in previous research (Wang, et al. 2006), this rich dataset of retrogenes and their rapid origination provided an opportunity to investigate and understand the evolutionary patterns of the retrogene pairs in rice, and to check whether the chimerical gene undergoes rapid positive selection subsequence from retrogene formed.

## Materials and Methods

### Samples, Primers and Molecular Cloning

There are ten species and two subspecies included in our study, the seven species, *Oryza grandiglumis* (use *Grandi* for short), *Oryza longistaminata (Longi), Oryza alta (Alta), Oryza australiensis (Austra), Oryza rufipogon (Rufi), Oryza nivara (Nivara a* and *Nivara* b), *Oryza glaberrima (Glab)*, get from International Rice Research Institute (IRGC), the IRGC ACC ID is shown in Table S1. The other two species *O. punctate (YSD8*) and *O. officinalis* (OWR) were from Wang’s lab. And the two subspecies *Oryza sativa* ssp. *Indica (Indica*) and *Oryza sativa* ssp. *japonica* (*Japonica*) were used as reference genomes for that whose whole genome have been completely sequenced and treated as gold standard. Total genomic DNA was isolated from leaf using the Cetyl Ttrimethyl Ammonium Bromide (CTAB) method. The *YSDB* (BB genome) and *OWR* (CC genome) genomic DNA were obtained from Wang’s lab.

All primers were designed according to genome sequences of *Japonica* and *Indica* in Table S2 (the other 17 pairs are not shown). Since the extremely redundant sequences around the chimerical retrogenes region, the primers were annealing to flanked sequence with approximate 1 kb length of PCR products. After amplified by polymerase chain reaction (PCR), the product DNA was sequenced with single-end from the 5’ends methods on an ABI Prism 3730 sequencer. All the sequence used in our study were derived from PCR sequencing unless PCR did not success in reference species but succeed in other sibling species. For this instance, substitution of 9311 genomic sequence was used for Indica in later analysis.

### Sequencing region detail

In previous study (Wang, et al. 2006), 898 intact retrogenes were found in *Indica* (9311) by *in-silico* way, and they indicated that 380 retrogenes have chimerical structures. We chose 24 retrogenes randomly from the 380 retrogenes, and positive selection acted on some specific branch (the analysis is show in the latter chapter) of seven retrogenes. The seven retrogenes are *RCG1* (Retro-Chimerical Gene1, chimerical id Chr03_4107, chimerical id is identical with the data in 2006 paper), *RCG2* (Chr04_4524), *RCG3* (Chr12_934), *RCG4* (Chr10_2602), *RCG5* (Chr01_5436), *RCG6* (Chr02_1920), *RCG7* (Chr08_3454). To exclude the artefacts of genome sequencing and assembly in 9311, we searched these seven chimerical retrogene and parental gene against newly PacBio genome IR8 (Table S3). According to the previous study and public database (Gramene), all these seven genes didn’t find homologous structure in maize and sorghum. The chimerical structure of three retrogenes are demonstrated in Fig. S3.

### Sequence edit and blast analysis

Using the designed primers, we cloned the sequences from the wild rice genomic DNA. The sequences got from the PCR were shown in Table S4. In the computational evolutionary analysis, the sequences cloned by PCR which is not long enough or can’t alignment to the retrogene is eliminated. In *RCG4* and *RCG7*, the *Indica* sequence from PCR (Indica in Fig. 1) share high similarity with reference genome (Indica_genome in Fig. 1), and we cannot confirm which one is orthologous to other species, so both PCR sequences and genomic sequences are used in the calculation for this study.

**Figure 1.**
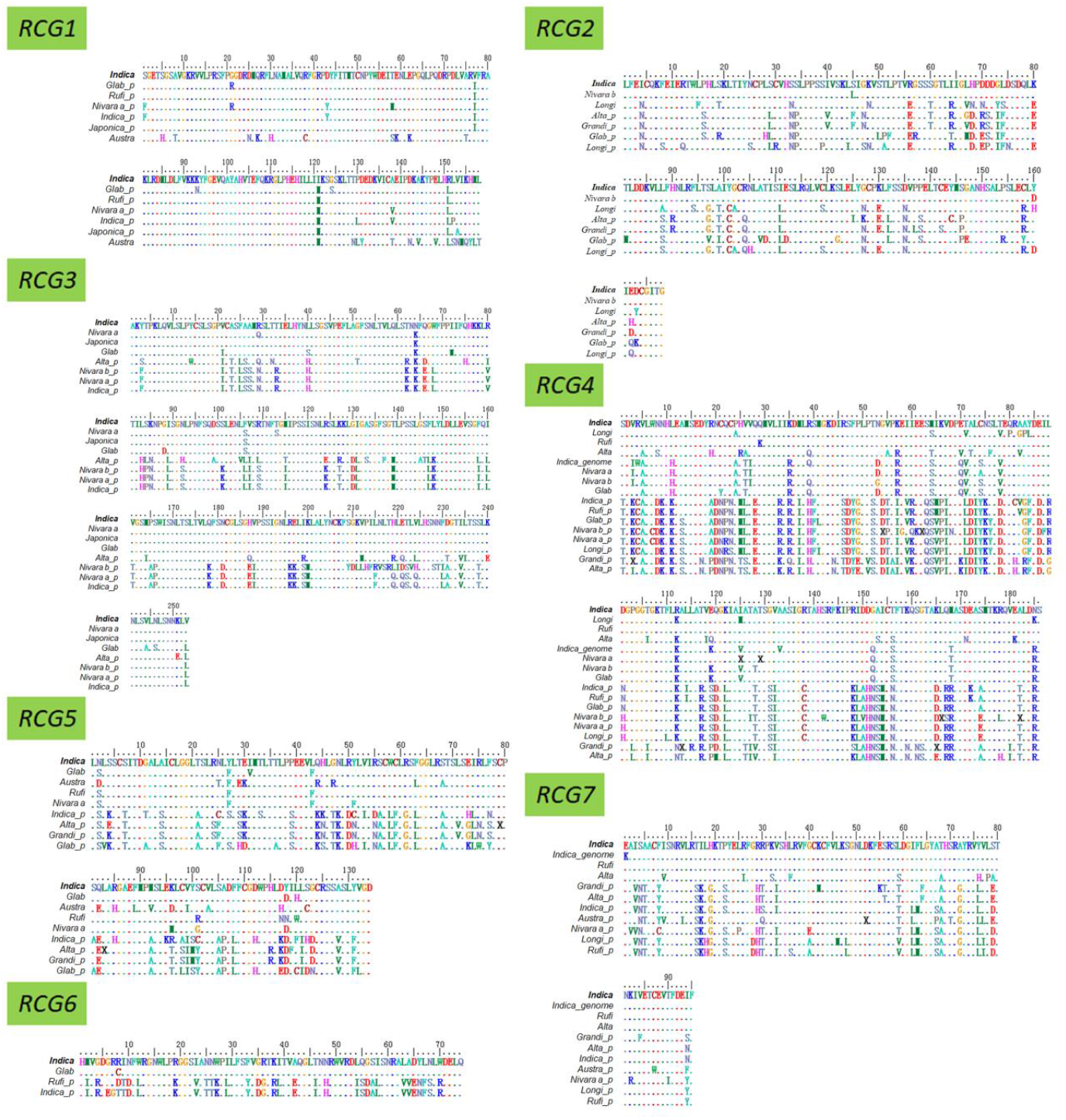
The amino acid alignment of seven chimerical retrogene pairs. *_p* represented the sequence of parental gene; *_genome* means the corresponding genomic region of Indica (9311) were used as substitutions of RCGs if it successfully amplified by PCR in sibling species but failed in 9311 or is differed from 9311 PCR results. Dot signify the amino acid was the same with that of 9311 in the alignment. The names of species are consistent with the short name of Table S1.

### Molecular evolution analysis

#### Phylogenetic Reconstruction

The sequences of retrogene pairs of coding regions were first translated to amino acid using the chimerical retrogene structure according to reference sequences, after the alignment by MEGA7 (Tamura, et al. 2007) with ClustalW, the amino acid sequences were retranslated into nucleotide. The amino acid alignments of seven positive selection candidate retrogene pairs were shown in Fig. 1, the other seventeen are shown in Fig. S1. The phylogenies used in analysis are built by MEGA7 using NJ method with the default parameter. All seven phylogenies were shown in Fig. 2, the other seventeen were shown in Fig. S2.

**Figure 2.**
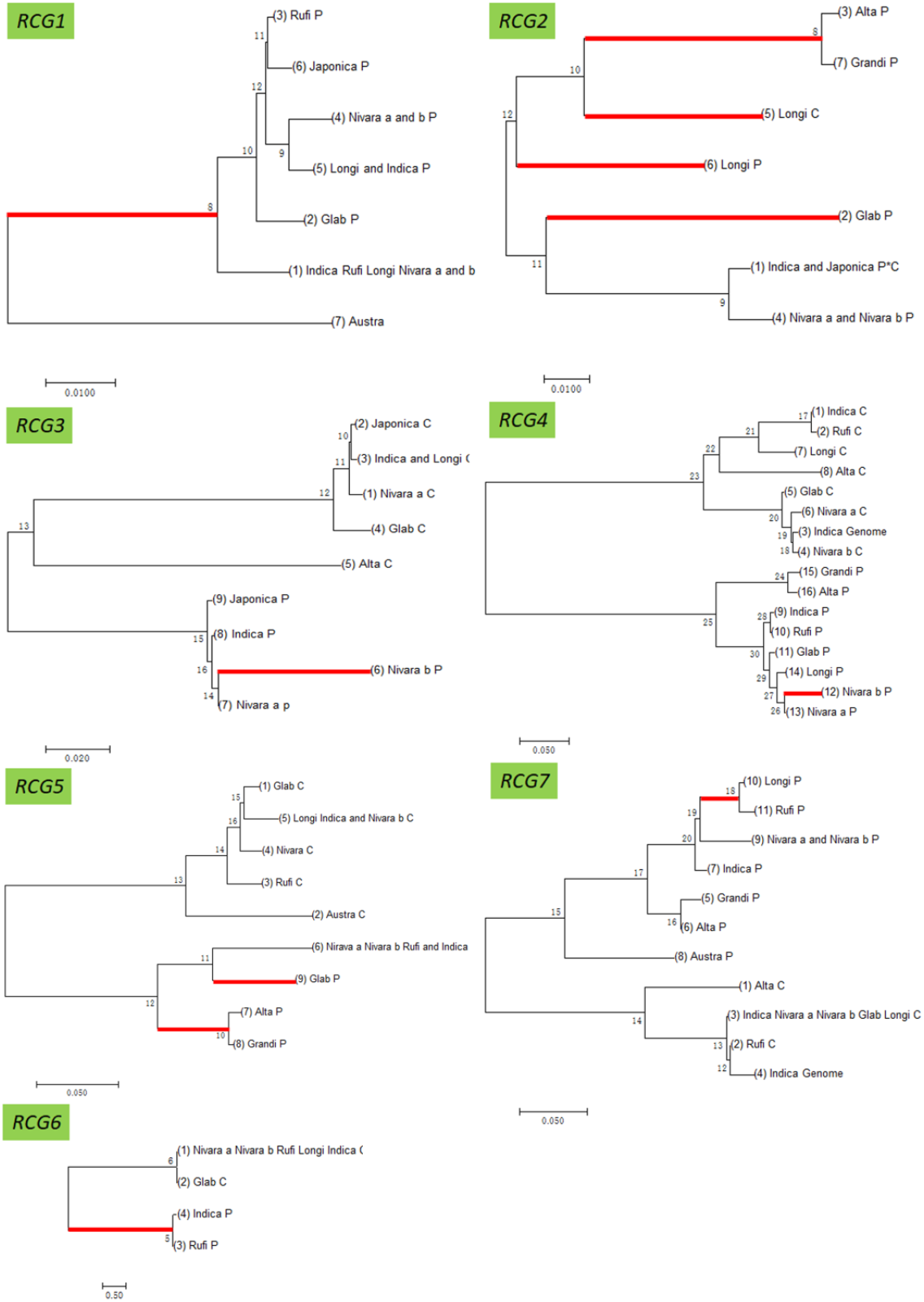
The phylogeny of seven chimerical retrogenes pairs. Phylogenetic tree was built in MEGA7 with default parameters. _p represented the sequence of parental gene and C means the chimerical retrogene; Genome suffixed in the specie name means the corresponding genomic region of Indica (9311) were used as substitutions of RCGs if it successfully amplified by PCR in sibling species but failed in 9311. Positive selection happened on the red bold branch. The species names are consistent with the shorted specie names of Table S1, & in the specie name represent the concatenated species share the identical sequence. The same for Fig. S2.

#### Maximum likelihood analysis for estimating the parameters

We employed the OBSM (Optimal Branch Specific Model) program (Zhang, et al. 2011) to explore the most probable branch-specific model to estimate its non-synonymous substitution per non-synonymous site (Ka) and synonymous substitution per synonymous site (Ks) respectively and the corresponding omega (ω = Ka/Ks) ratio. Here, ω is well accepted in evolutionary interpretation that when ω>1, suggesting positive selection; when ω≈1, suggesting neutral evolution while ω<1 suggest purify selection with functional constraint. OBSM has three methods, the first method cost less time while the third method is more time-consuming but gets a better result, which means that have a better branch-specific model in likelihood ratio test (LRT) (Zhang, et al. 2011) or Akaike Information Criterion (AIC) comparison (Akaike 1974).

We calculated all these 24 retrogene sets by three methods of OBSM. In analysis, we removed all gaps in alignments, set the codon frequency of the CODEML control file at CodonFreq = 3, set the parameter k in method III of OBSM at 0.5. Furthermore, we employed the branch-site model (Yang and Nielsen 2002) to explore the positive sites, and fix the specific branch suggested by the final optimal models as foreground branch. The suggested test 1 and the suggested test 2 were employed to detect positive selection sites (Zhang, Nielsen, et al. 2005).

## Results

### Seven retrogene pairs undergo positive selection

According to the results of calculation by three methods, we obtain seven among twenty-four retrogene pairs were undergoing positive selection. All the log likelihood (lnL) values and parameter of final optimal models for seven retrogene pairs for each method are shown in Table 1; other seventeen retrogenes are shown in Table S5, which laid foundations for the selective site analysis in Table 2. All these analyses are described in detail as follows.

**Table 1.**
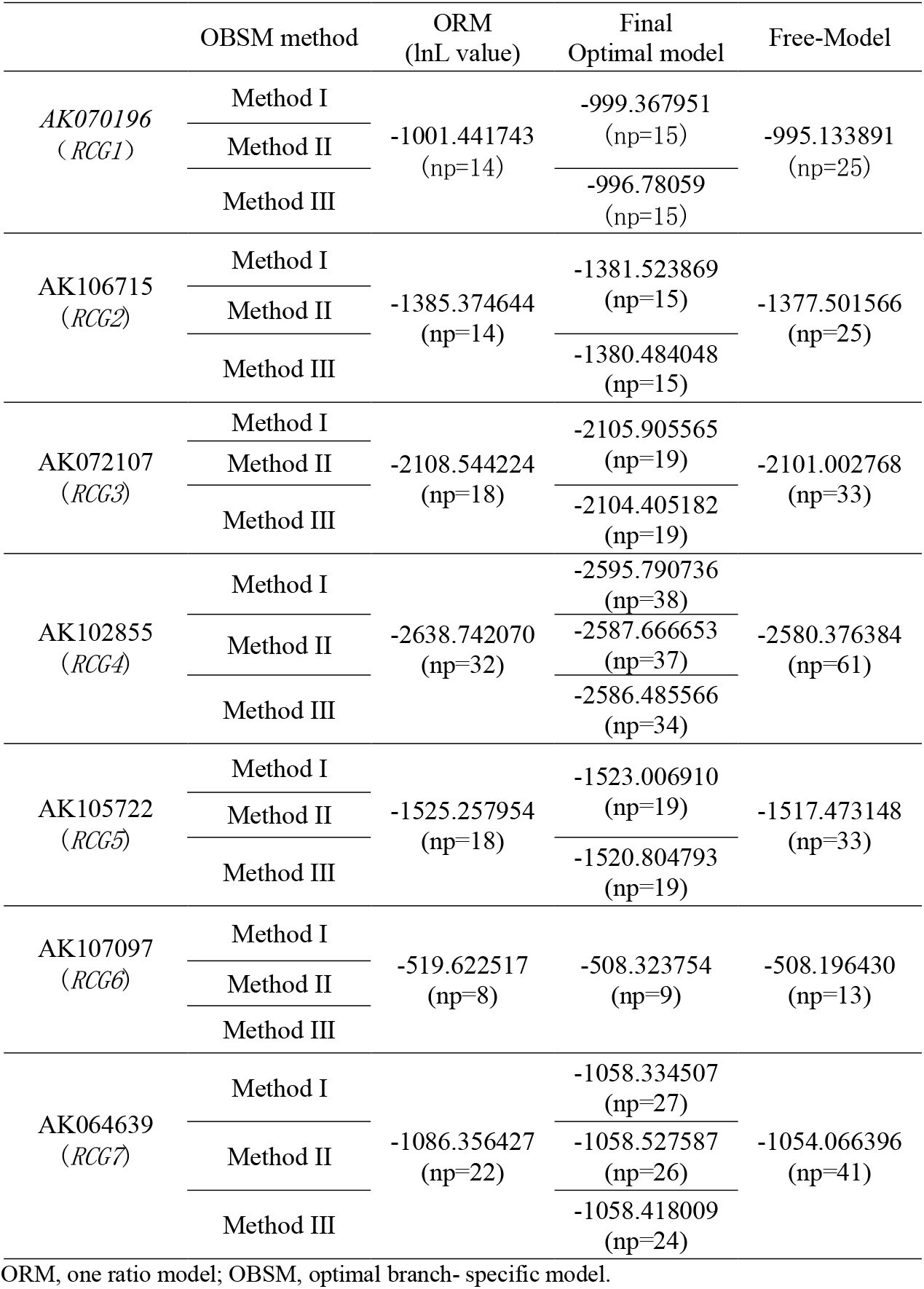
Log likelihood value of seven chimerical retrogene pairs.

**Table 2.**
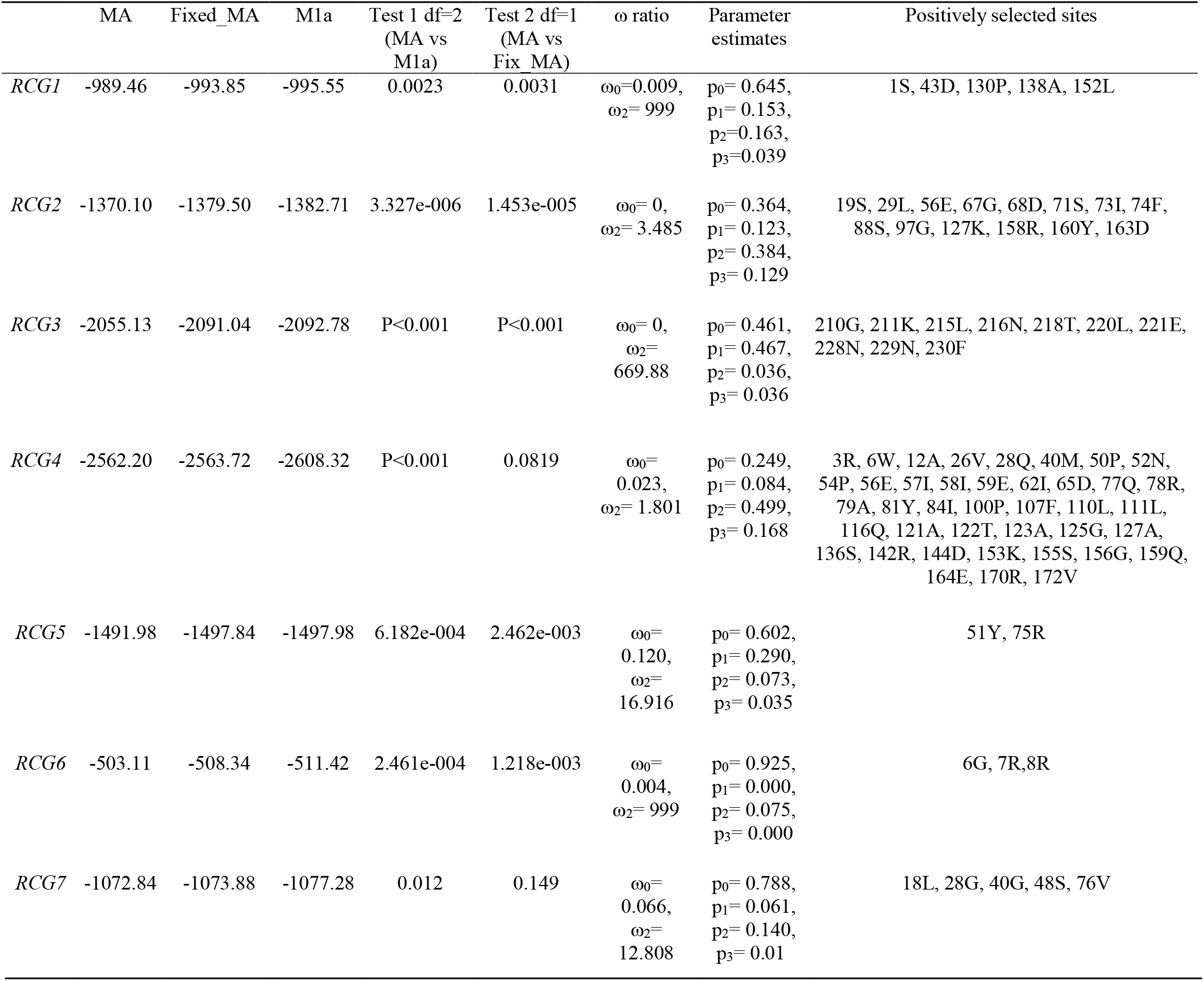
Branch-site method estimation of seven chimerical retrogene pairs. MA, model A of branch-site model analysis in PAML.

| | MA | Fixed_MA | M1a | Test 1 df=2<br>(MA vs<br>M1a) | Test 2 df=1<br>(MA vs<br>Fix_MA) | $\omega$ ratio | Parameter<br>estimates | Positively selected sites |
| --- | --- | --- | --- | --- | --- | --- | --- | --- |
| *RCG1* | -989.46 | -993.85 | -995.55 | 0.0023 | 0.0031 | $\omega_0=0.009$,<br>$\omega_2=999$ | $p_0=0.645$,<br>$p_1=0.153$,<br>$p_2=0.163$,<br>$p_3=0.039$ | 1S, 43D, 130P, 138A, 152L |
| *RCG2* | -1370.10 | -1379.50 | -1382.71 | 3.327e-006 | 1.453e-005 | $\omega_0=0$,<br>$\omega_2=3.485$ | $p_0=0.364$,<br>$p_1=0.123$,<br>$p_2=0.384$,<br>$p_3=0.129$ | 19S, 29L, 56E, 67G, 68D, 71S, 73I, 74F,<br>88S, 97G, 127K, 158R, 160Y, 163D |
| *RCG3* | -2055.13 | -2091.04 | -2092.78 | P<0.001 | P<0.001 | $\omega_0=0$,<br>$\omega_2=669.88$ | $p_0=0.461$,<br>$p_1=0.467$,<br>$p_2=0.036$,<br>$p_3=0.036$ | 210G, 211K, 215L, 216N, 218T, 220L, 221E,<br>228N, 229N, 230F |
| *RCG4* | -2562.20 | -2563.72 | -2608.32 | P<0.001 | 0.0819 | $\omega_0=0.023$,<br>$\omega_2=1.801$ | $p_0=0.249$,<br>$p_1=0.084$,<br>$p_2=0.499$,<br>$p_3=0.168$ | 3R, 6W, 12A, 26V, 28Q, 40M, 50P, 52N,<br>54P, 56E, 57I, 58I, 59E, 62I, 65D, 77Q, 78R,<br>79A, 81Y, 84I, 100P, 107F, 110L, 111L,<br>116Q, 121A, 122T, 123A, 125G, 127A,<br>136S, 142R, 144D, 153K, 155S, 156G, 159Q,<br>164E, 170R, 172V |
| *RCG5* | -1491.98 | -1497.84 | -1497.98 | 6.182e-004 | 2.462e-003 | $\omega_0=$<br>0.120,<br>$\omega_2=$<br>16.916 | $p_0= 0.602,$<br>$p_1= 0.290,$<br>$p_2= 0.073,$<br>$p_3= 0.035$ | 51Y, 75R |
| *RCG6* | -503.11 | -508.34 | -511.42 | 2.461e-004 | 1.218e-003 | $\omega_0=$<br>0.004,<br>$\omega_2= 999$ | $p_0= 0.925,$<br>$p_1= 0.000,$<br>$p_2= 0.075,$<br>$p_3= 0.000$ | 6G, 7R,8R |
| *RCG7* | -1072.84 | -1073.88 | -1077.28 | 0.012 | 0.149 | $\omega_0=$<br>0.066,<br>$\omega_2=$<br>12.808 | $p_0= 0.788,$<br>$p_1= 0.061,$<br>$p_2= 0.140,$<br>$p_3= 0.01$ | 18L, 28G, 40G, 48S, 76V |

#### RCG1

*RCG1* is a new gene that originated 3.15 MYA (Ks≈0.041) in the rice genome. The log likelihood (lnL) value of the optimal model of method III is −996.78, is significantly better than the lnL value of the optimal model of the method I and method II (LRT: df =1 2ΔL=5.17 p-value=0.023). This result indicates that method III more suitable for *RCG1* data. The estimating of Ka/Ks ratio of lineage branch 9 in the final optimal model of the method I and method II were infinite (999), and the Ka/Ks ratio of branch 9, 8, 11 and 5 in the final optimal model of method III is infinite (999). All these models indicate that the evolution pattern of *RCG1* retrogene pair is episodic. Although it failed in likelihood ratio test (LRT: df =1, 2ΔL=3.006, p-value=0.083) when we nested a comparison between the final optimal model and fix-model which fixed the Ka/Ks ratio of branch 9, 8, 11 and 5 to one, the estimates of parameters in this optimal model suggest that there’re sixteen non-synonymous substitutions versus zero synonymous substitution occurred along the lineage 8, it has a great possibility that the lineage 8 is undergoing positive selection that the previous study suggest positive selection when the non-synonymous substitutions greater than 9 while the synonymous substitution is equal to 0 (Nozawa et al. 2009). Based on the final optimal model of method III, we used the branch-site model to identify the positive sites. In test 1, M1a (lnL=-995.55) versus Model A (lnL=-989.46), 2Δl= 12.17, p-value=0.0023 (df=2); in test 2, Model A versus fix-Model A (lnL=-993.85), 2Δl= 8.77, p-value=0.0031 (df=1). All these two tests indicate that the Model A fit the data better than others, Model A suggests five sites to be potentially under positive selection along the foreground branch at the 95% level according to the BEB analysis, these sites are 1S, 43D, 130P, 138A, 152L, the parameters estimate by Model A are p0= 0.645, p1= 0.153, p2= 0.163, p3=0.039, ω0=0.009, ω2= 999.

#### RCG2

*RCG2* is a new gene that originated 6.92 MYA (Ks≈0.090) in the rice genome. The OBSM methods suggest that, excepting lineage 4 in final optimal model of the method I and method II, lineage 4 *Nivara* a and b_P and lineage 1 *Indica-Japonica* P&C in final optimal model of the method III, the Ka/Ks ratio is less than 1 (0.358, 0.321 respectively), all other lineages are greater than 1 (1.744, 1.835 respectively). The log likelihood (lnL) values of these two models are −1381.52 and −1380.48, respectively. Since they have the same ω ratio numbers, the latter model is considered being better because of lower lnL value. The *RCG2* retrogene pair were undergoing positive selection is confirmed when we nested a comparison between the fix-model and corresponding final optimal models, the 2ΔL is 6.474, the p-value is 0.011. The final optimal model indicates that the positive selection permeates the whole evolution pattern of *RCG2* retrogene pair. The estimates of parameters in the final optimal models suggest that the non-synonymous substitutions in five lineages 3, 7, 5, 6 and 2 are all greater than 9, rang from 10.5 to 26.3.

Model A more suitable than others based on the final optimal model, two branch-sites model tests. Nine sites to be potentially under positive selection along the foreground branch at the 95% level according to the BEB analysis (19S, 29L, 56E, 67G, 68D, 71S, 73I, 74F, 88S, 97G, 127K, 158R, 160Y, 163D). The parameters suggested by Model A are p0= 0.364, p1= 0.123, p2= 0.384, p3= 0.129, ω0= 0, ω2= 3.485.

#### RCG3

*RCG3* is homologous to a *Verticillium wilt* resistance gene *Ve1* (Kawchuk, et al. 2001; Fradin, et al. 2009) which originated 14.77 MYA (Ks≈0.192) in the rice genome. The lnL value of final optimal model of the method I and method II is −2105.91, the lnL value of the final optimal model of method III is −2104.41, since they have the same ω ratio numbers, the latter model is considered being better. The estimate of Ka/Ks ratio of lineage *Nivara* b_P in final optimal model of the method I and method II is 1.388, the estimate of Ka/Ks ratio of branch 15, 6 and 10 in the final optimal model of method III is 1.524. Although all these two models not significant in LRTs tests when we nested a comparison between the fix-model and final optimal model, it is suggested that the branch 8 have a much higher substitution rate than the background substitution rate since the large non-synonymous substitutions in it (30.3 and 31.0 respectively).

Based on the final optimal model, two branch-sites model tests based on the final optimal models indicate that the Model A fit the data better than others. Model A suggests ten sites to be potentially under positive selection along the foreground branch at the 95% level according to the BEB analysis, these sites are 210G, 211K, 215L, 216N, 218T, 220L, 221E, 228N, 229N, 230F. Surprisingly, all these sites are very close to each other and seem to be a functional domain. The parameters suggested by Model A are p0= 0.461, p1= 0.467, p2= 0.036, p3= 0.036, ω0= 0, ω2= 669.88.

#### RCG4

Given the complexity of these sixteen sequences included in this retrogene pair, the result of the most probable estimating models suggested by OBSM are different totally. The final optimal model suggested by Method I is a seven-ratio model and the lnL value is −2595.79. The final optimal model suggested by Method II is a six-ratio model and the lnL value is −2587.67. The final optimal model suggested by Method III is a three-ratio model and the lnL value is −2586.49. Obviously, the final optimal model of Method III fit the data better than other two models since the fewer parameters and the larger lnL value. Although this model failed in LRTs when we nested a comparison between the fix-model and final optimal model, it is suggested by all three final optimal models that the lineage *Nivara* b_P have a much higher substitution rate than the background substitution rate. The estimates of parameters in these three optimal models suggest that the non-synonymous substitutions in lineage *Nivara* b_P are 18.7, 18.7 and 16.5 respectively.

Based on the final optimal model of method III, two tests indicate that the Model A fit the data better than other models. Model A suggests two sites to be potentially under positive selection along the foreground branch at the 95% level according to the BEB analysis; these sites are 51Y, 75R. The parameters suggested by Model A are p0= 0.602, p1= 0.290, p2= 0.073, p3= 0.035, ω0= 0.121, ω2= 16.92.

#### RCG5

The lnL value of the final optimal model of Method I and Method II is −1523.01, the lnL value of final optimal model of Method III is −1520.80, the latter one is significantly better than the former one according to the LRTs (df=1, 2ΔL=4.404, p-value=0.036). This result indicates that the final optimal model of method III fit *RCG5* gene pair better than the former model. The estimating of Ka/Ks ratio of lineage Glab_P in final optimal model of method I and method II is 2.20, and the estimating of Ka/Ks ratio of lineage Glab_P, branch 10, and lineage *Nivara* a in the final optimal model of method III is 2.66. All these models indicate that the evolution pattern of RCG5 retrogene pair is episodic. Although it is failed in LRTs (df=1, 2ΔL=2.612, p-value=0.106) when we nested a comparison between the final optimal model and fix-model which fixed the Ka/Ks ratio of lineages Glab_P, branch 10 and Nivara-a equals to one. The estimates of parameters in final optimal model of method III suggest that they’re about 10.8 non-synonymous substitutions along the branch 10, and there’re 16.6 non-synonymous substitutions along the lineage Glab_P, it has a great possibility that the branch 10 and Glab_P are undergoing positive selection.

Based on the final optimal model of method III, we used branch-site model to identify the positive sites. In test 1, M1a (lnL=−995.55) versus Model A (lnL=−989.46), 2Δl= 12.172, p-value=0.0023 (df=2), in test 2, Model A versus fix-Model A (lnL=-993.85), 2Δl= 8.770, p-value=0.0031 (df=1). All these two tests indicate that the Model A fits the data better than others, Model A suggests five sites to be potentially under positive selection along the foreground branch at the 95% level according the BEB analysis, these sites are 1S, 43D, 130P, 138A, 152L, the parameters suggested by Model A are p0= 0.645, p1= 0.153, p2= 0.163, p3=0.0387, ω0=0.00935, ω2= 999.

#### RCG6

The three OBSM methods suggested an identical final optimal model. The estimating of Ka/Ks ratio except branch 5 is suggested to be infinite (999). Although it is failed in LRTs (df =1 2ΔL=3.108 p-value=0.0779) when we nested a comparison between the final optimal model and fix-model which fixed the Ka/Ks ratio of all lineages equal to one except branch 5, the estimates of parameters in this optimal model suggest that they’re about 19.5 non-synonymous substitutions versus 7.1 synonymous substitutions occurred along the branch 10, it has a great possibility that the lineage B is undergoing positive selection.

Based on the final optimal model, we used branch-site model to identify the positive sites. In test 1, M1a (lnL=-511.42) versus Model A (lnL=-503.11), 2Δl= 16.62, p-value=2.461e-004 (df=2), in test 2, Model A versus fix-Model A (lnL=-508.34), 2Δl= 10.46, p-value=1.218e-003 (df=1). All these two tests indicate that the Model A fit the data better than others, Model A suggests three sites to be potentially under positive selection along the foreground branch at the 95% level according to BEB analysis, these sites are 6G, 7R, 8R, the parameters suggested by Model A are p0= 0.925, p1= 0.00, p2= 0.0753, p3= 0.00, ω0= 0.0045, ω2= 999.

#### RCG7

Given the complexity of these eleven sequences included in this retrogene pair, the result of the most probable estimating models suggested by OBSM are all different. The final optimal model suggested by Method I is a six-ratio model and the lnL value is −1058.33. The final optimal model suggested by Method II is a five-ratio model and the lnL value is −1058.53. The final optimal model suggested by Method III is three-ratio model and the lnL value is −1058.42. Although the final optimal model of the Method III has fewer parameters than other two models, the lnL value of these three models are very close to each other. This final optimal model of Method III suggested the Ka/Ks ratios of all lineages are less than one while other two models all suggested the branch 18 and lineage *Grandi_P* are larger than one. Although all LRTs comparisons between the final optimal models of Method I and Method II and fix-model in which fix branch 18 and lineage *Grandi_P* equal to one are failed, it is suggested by two final optimal models that the branch 18 have a much higher substitution rate than the background substitution rate since the estimates of parameters suggest that there’re 7.6 non-synonymous substitutions versus 1.1 synonymous substitutions occurred along the branch 18.

We used the branch-site model to identify the positive sites, the suggested test 1 and the suggested test 2 are employed to detecting positive selection sites along branch 18. Test 1 suggested that Model A is significantly better than the model M1a while it is failed in test 2. Model A suggests five sites to be potentially under positive selection along the foreground branch at the 95% level according to the BEB analysis; these sites are 18L, 28G, 40G, 48S, 76V. The parameters suggested by Model A are p0= 0.788, p1= 0.0612, p2= 0.140, p3= 0.0109, ω0= 0.0662, ω2= 12.81.

### Tajima’ D test suggests the mutations in RCG4, RCG6 are deviation from neutral mutation hypothesis

Whether retrogenes under neutral selection? We also employed Tajima’ D test included in MEGA 7 to check the mutations in chimerical retrogene (Tajima 1989). The result suggested only chimerical retrogene *RCG4* and *RCG6* pair are significant, while the mutations among the other four retrogene pairs are deviation from neutral. The significant deviation of D from 0 is observed in *RCG4* (p<0.01) and in *RCG6* (p<0.001), the detail is shown in Table 3.

**Table 3.** Results of Tajima’s Neutrality Test for seven chimerical retrogene pairs.

|  | m | S | p<sub>s</sub> | Θ | π | D |
| --- | --- | --- | --- | --- | --- | --- |
| *RCG4* | 16 | 313 | 0.570 | 0.172 | 0.270 | 2.486 |
| *RCG6* | 4 | 79 | 0.357 | 0.195 | 0.240 | 2.443 |

The Tajima test statistic was estimated using MEGA7. All positions containing gaps and missing data were eliminated from the dataset (Complete deletion option). The abbreviations used are as follows: m = number of sites, S = Number of segregating sites, p<sub>s</sub> = S/m, Θ = p<sub>s</sub>/a1, and π = nucleotide diversity. D is the Tajima test statistic.

### The patterns of substitutions in new retrogenes and parental genes

Three distinct patterns have been revealed base on synonymous and replacement sites in the seven gene pairs were shown in Fig. 2. (1) The chimerical genes were rapidly substituted in the initial stage of the new gene lineage under positive selection, e.g. *RCG2*. This is partially consistent with the pattern revealed by Jones and Begun (Jones and Begun 2005; Jones et al. 2005), three *Adh* related new retrogenes evolved rapidly after the new gene were formed. Furthermore, our result suggests the rapid evolution also happened to parental gene. This type of rerouted functional evolution covered several occasions: (2) The parental genes evolved rapidly soon after the chimerical genes were formed whereas the new genes evolved slowly in evolution. *RCG6* belongs to this category. (3) The parental genes evolved after some uncertain period of the chimerical genes were formed whereas the new genes evolved slowly in evolution, shown as *RCG3, RCG4, RCG5* and *RCG7*. Both pattern (2) and (3) implicated an unexpected process of evolution in functionality: the new retrogenes might replace the parental gene to carry out the ancestral functions while the parental gene might have evolved new functions driven in adaptive evolution.

### *RCG3* may plays an important role in disease resistance

We compared our seven chimerical retrogenes to the probesets of Rice Genome Arrays of Affymetrix GeneChip, since the high complexity and the redundancy of the retro gene similar copy (Table 5) and the incomplete probesets coverage of rice genome, only pairs of *RCG3* and *RCG5* have the perfect match probesets, the compared detail is shown in Table 4, the expression profile can be got from the CERP database (http://crep.ncpgr.cn/). However, both *RCG3* and *RCG5* showed functional divergence (Fig. 3). Especially, according to the entire life cycle of rice gene expression data (Wang, et al. 2010), chimerical retrogene *RCG3* probe (Os.54355.1.S1_at) has an expression peak in Zhenshan 97 (a variety of cultivated rice) at infection period in calli, germination period (72h after imbibition) in seed and 21 days after pollination in endosperm. This result is in consonance with the independent evidence from the TIGR (http://rice.plantbiology.msu.edu) that this gene encodes Leucine-rich proteins, and has a high similarity with the *Ve1* gene which has been shown to be resistant to *Verticillium wilt* disease (Fradin et al. 2009; Kawchuk et al. 2001).

**Figure 3.**
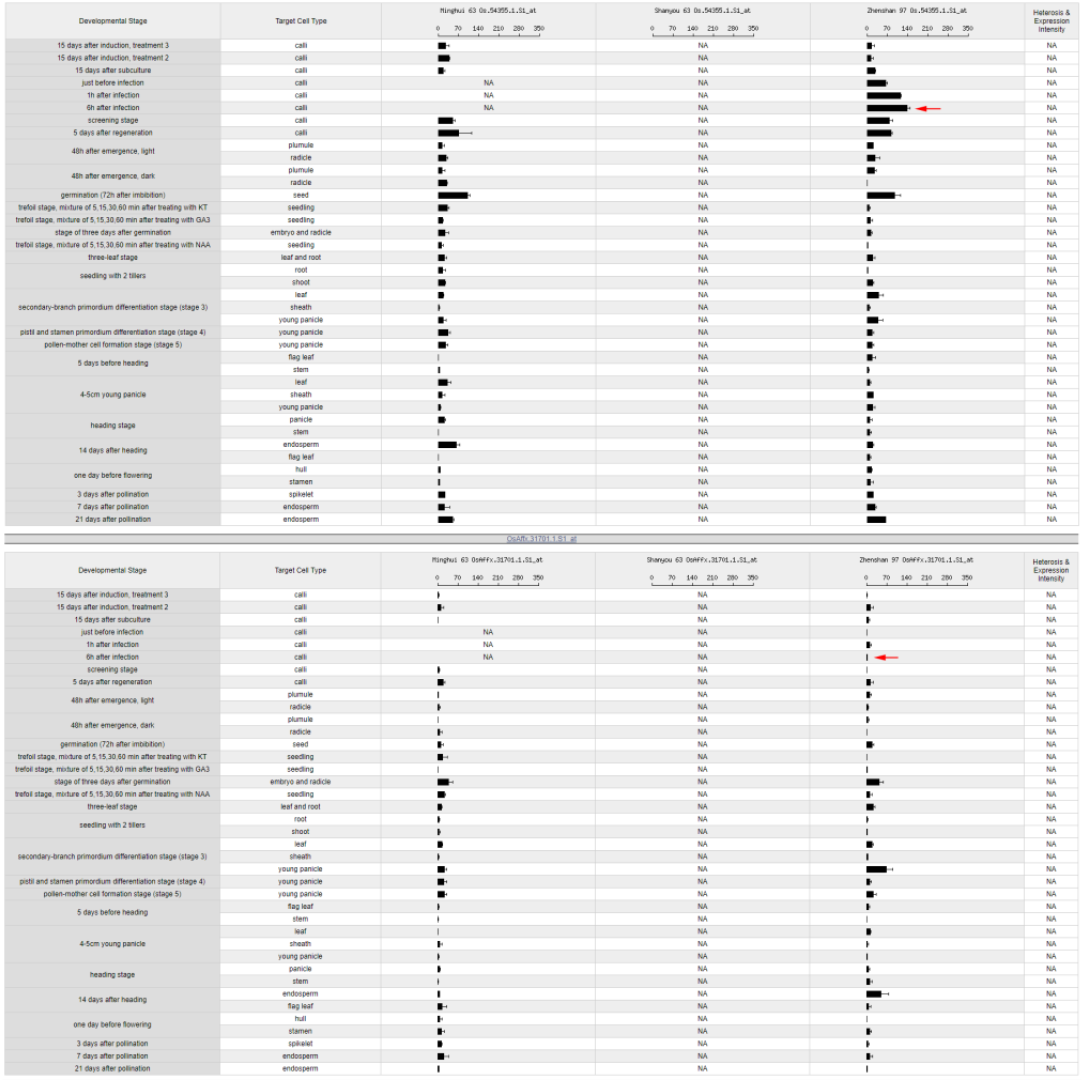
Divergent expressions between *RCG3* and its parental gene. The corresponding sequences were searched against Affymetrix Rice Genome Array, and the digital expression profiles were generated automatically in http://crep.ncpgr.cn/crep-cgi/blast.pl. Red arrow for chimerical retrogene indicate the highest expression stage, however, red arrow for parental gene point to the expression of same stage.

**Table 4.** Affymetrix GeneChip expression profile of seven chimerical retrogene pairs.

|  | Chimerical retrogene ID in Plant cell paper | Chimerical Affy Probset names | Parental Affy Probset names |
| --- | --- | --- | --- |
| *RCG1* | Chr03_4107, AK070196_Chr03_27608263_27613159 | NA | NA |
| *RCG2* | Chr04_4524, updata_AK106715_Chr04_30664045_30669070 | Os.57563.1.S1_at | NA |
| *RCG3* | Chr12_904, updata_AK072107_Chr12_5820378_5826726 | Os.54355.1.S1_at | OsAffx.31701.1.S1_at |
| *RCG4* | Chr10_2602, updata_AK102855_Chr10_17747411_17752061 | NA | OsAffx.29724.1.S1_at |
| *RCG5* | Chr01_5436, updata_AK105722_Chr01_36521616_36526443 | Os.35231.1.S1_at | Os.50239.1.S1_a_at |
| *RCG6* | Chr02_1920, updata_AK107097_Chr02_12785386_12789823 | NA | Os.54261.S1_at |
| *RCG7* | Chr08_3454, updata_AK064639_Chr08_24470676_24475311 | NA | NA |

Sequences of chimerical gene and its parental gene were searched against rice expression profile CREP (http://crep.ncpgr.cn/crep-cgi/home.pl). Probe applied to target sequence only when no mismatch (e-value=0) and hybrid to the right position. NA, no perfect match was found for chimerical retrogene pairs.

**Table 5.** Copy number variation of similarity hits in OMAP/OGE genomes.

|  | RCG1 | RCG2 | RCG3 | RCG4 | RCG5 | RCG6 | RCG7 | Genome_size (Mb) |
| --- | --- | --- | --- | --- | --- | --- | --- | --- |
| *Oryza barthii* | 118 | 106 | 12000 | 59 | 104 | 39 | 794 | 760 |
| *Oryza brachyantha* | 10 | 55 | 6667 | 47 | 28 | 4 | 729 | 389 |
| *Oryza glaberrima* | 131 | 102 | 11609 | 60 | 90 | 30 | 1240 | 389 |
| *Oryza longistaminata* | 89 | 214 | 806 | 129 | 95 | 48 | 217 | 760 |
| *Oryza meridionalis* | 161 | 36 | 11201 | 70 | 80 | 34 | 298 | 760 |
| *Oryza nivara* | 136 | 54 | 12000 | 81 | 90 | 36 | 821 | 539 |
| *Oryza punctata* | 1148 | 38 | 9116 | 96 | 58 | 13 | 1776 | 1691 |
| *Oryza rufipogon* | 160 | 79 | 12000 | 120 | 122 | 37 | 1315 | 1201 |
| *Oryza sativa indica* | 146 | 84 | 12107 | 155 | 124 | 29 | 1382 | 1000 |
| *Oryza sativa japonica* | 142 | 67 | 12037 | 158 | 125 | 35 | 1678 | 1054 |

The genomic sequences of seven RCGs were blastn searched against Gramene database with the e-value threshold of 1e-5.

### Chimerical retrogene *RCG1* is a young gene

The Ks value for seven retrogenes was calculated based on the simple two sequences (parental verse new genes) comparison (Wang et al. 2006). The values were 0.124, 0.19, 0.281, 2.27, 0.547, 1.884 and 3.575. Because more sequences data have been available, we recalculated the Ks value for *RCG1, RCG2* and *RCG3* by MEGA7 using the NG86 model (Nei and Gojobori 1986; Zhang, et al. 1998) with the transition/transversion ratio k=2. To estimate the divergence time accurately, since the branch *Austra* (Fig. 2) is ancestral to the clade generated by the retroposition event, this branch is excluded from *RCG1* data in the analysis. Then the Ks values with the 95% confidence interval for *RCG1, RCG2* and *RCG3* are 0.041±0.011, 0.090±0.016 and 0.192±0.021 respectively. Assuming that the synonymous substitution rate of rice genes is 6.5 × 10^−9^ substitutions per site per year (Gaut, et al. 1996), then these chimerical retrogenes would have been formed around 3.15±0.88 MYA, 6.92±1.23 MYA and 14.77±1.62 MYA. These estimates suggest that the three chimerical retrogenes are very young (*RCG1* and *RCG2*) or young (*RCG3*).

## Discussion

In this study, we used the program OBSM (Zhang et al. 2011) to explore the optimal branch model for chimerical genes. OBSM is CODEML (one program included in PAML package) (Yang 2007) aid programs which help the user to found out the optimal branch-specific models (Yang 1998) using the maximum likelihood approach. We also used the branch-site approach to explore positive selection sites, although we note this method have some defects like it may not suggest right sites proposed by Nozawa, et al. (2009). In fact, in our data analysis, especially in *RCG3*, the sites suggested by MA model seem reasonable; because these sites are all belong to Leucine-rich repeat region which may have some connection with disease resistance. The disease resistance function may help the individual with better adaption to be selected to survive.

The common patterns and mechanisms shaping the evolution of new genes were generalized by many previous studies. Corbin D. Jones (Jones and Begun 2005; Jones et al. 2005) analyzed the origination of three *Drosophila* gene *jinwei, Adh-Finnegan*, and *Adh-Twain*, and unveiled three genes underwent rapid adaptive amino acid evolution in a short time after they were formed, followed by later quiescence and functional constraint. In 2008, study of novel alcohol dehydrogenase *siren1* and *siren2* also proved that chimerical genes evolved adaptively shortly after they were formed (Shih and Jones 2008). However, our results seem to indicate another different pattern, that is, besides the rapid adaptive amino acid evolution happened shortly after chimerical retrogene were formed, the rapid adaptive evolution also appeared in parental genes. This quickly evolution of parental gene occupied a high proportion in our seven chimerical retrogene pairs, six (*RCG2* to *RCG7*) of which have rapid adaptive evolution in parental gene evolution. The difference between *Drosophila* and *Oryza* may be caused by high proportion of retrotransposon in rice (McCarthy, et al. 2002; Baucom, et al. 2009; Paterson, et al. 2009), or because of the polyploidy origin of the rice genome and additional a recent segmental duplication occurred c. 5 MYA (Wang, et al. 2005). Subsequent large-scale chromosomal rearrangements and deletions may play an impact on the evolution pattern of chimerical retrogene pairs.

To compare the expression profile of *RCG3* and its parental gene, we locate the *RCG3* parental gene in *Japonica* genome and the located region is predicted as loci LocOs12g11370 by TIGR. The probeset (OsAffx.31701.1.S1_at) in this region reveal that the parental gene has an expression peak at secondary-branch primordium differentiation stage (stage 3) at young panicle (Fig. 3), while its parental gene only showed negligible signal for this stage. This is reasonable because the high expression level at generative organ may capture a higher chance to retroposition among the genome sequences.

In our analysis, seven out of twenty-four (29.17%) chimerical retrogene pairs seem to be undergoing positive selection. This proportion is much higher than that of previous whole-genome research in *Streptococcus* (Anisimova, et al. 2007) and *Apis mellifera* (Zayed and Whitfield 2008). The phylogenomic analysis of *Streptococcus* (Anisimova, et al. 2007) shows that 136 gene clusters out of 1730 (7.86%) underwent positive selection. Genome-wide analysis of positive selection in honey bee suggested that positive selection acted on a minimum of 852–1,371 genes or around 10% of the bee’s coding genome (Zayed and Whitfield 2008). If we consider 10% coding genes of whole genome undertake positive selection as the average, then the proportion 29.17% of chimerical retrogene is significantly higher than the average in Fisher exact test (p=0.001). We speculated that reverse transcripted mRNA intermediated new chimerical retrogene pairs have advantages for survival or propagation.

## Acknowledgments

We thank Shiping Wang for valuable discussion and support. This research was financially supported by the National Natural Science Foundation of China (grant number 31571311), the CAS “Light of West China” Program (grant number 292017312D11022), and partly supported by the open funds of the National Key Laboratory of Crop Genetic Improvement (grant number ZK201605).

## Supporting Information

Fig. S1 The amino acid alignment of seventeen chimerical retrogene pairs.

Fig. S2 The phylogeny of seventeen chimerical retrogene pairs.

Fig. S3 Paradigm of the chimerical retrogene model. Colorful rectangular boxes represent the exons, greyish boxes represent introns. Superordinate gene in each model is parental gene, lower part in each model is chimerical retrogene. Solid lines mean the border of homologous block and numbers designate the relative position.

Table S1 Species used in our analysis.

Table S2 Primers for PCR and sequencing.

Table S3 Chimerical retrogene and parental gene in IR8. The sequence of chimerical retrogene and corresponding parental gene were blat searched against Indica rice genome IR8, which was sequenced by Pacbio technology. Round brackets indicated the output of blat; angle brackets mean when blat out were too long, the sequences range were narrowed down by gene-specific primer.

Table S4 PCR based sequencing statistics of retrogenes and parental genes. C: Means the retro-chimerical gene; P: Means the parental gene; x: Means did not get PCR result; na: Means did not get valuable sequence; *: using the *Indica* reference sequence; &: The cloned sequence did not perfect match the reference sequence of 9311. Total sequences numbers, means the number of sequence type used for phylogeny construction, which correspond to the maximum value in C and P column for each retrogene.

The different number in the two columns of each retrogene represent a sequence type that unique for one or several species, which consistent with the sequence number of phylogenies in Fig.2.

Table S5. The lnL value comparison and the most probable model suggestion. Model fitting was optimized in OBSM (Zhang et al., 2011). *, significant at *p*<0.05; **, significant at *p*<0.01.

