## Supplemental tables for "Evolutionary patterns of the chimerical retrogenes in Oryza"

Table S1 Species used in our analysis

| ID | Species/subspecies | Short Name | Genome | IRGC ACC |
|---|---|---|---|---|
| 1 | *Oryza grandiglumis (Doell Prod.)* | *Grandi* | CCDD | 105664 |
| 2 | *Oryza longistaminata (A. Chev. et Roehr)* | *Longi* | AA | 103886 |
| 3 | *Oryza alta (Swallen)* | *Alta* | CCDD | 100967 |
| 4 | *Oryza australiensis (Domin)* | *Austra* | EE | 86530 |
| 5 | *Oryza rufipogon (Griff.)* | *Rufi* | AA | 80643 |
| 6 | *Oryza nivara (Sharma et Shastry)* | *Nivara a* | AA | 80622 |
| 7 | *Oryza nivara (Sharma et Shastry)* | *Nivara b* | AA | 80582 |
| 8 | *Oryza glaberrima (Steud.)* | *Glab* | AA | 103600 |
| 9 | *Oryza Sativa L. indica* | *Indica* | AA | / |
| 10 | *Oryza Sativa L. japonica* | *Japonica* | AA | / |
| 11 | *Oryza punctata* | *YSD8* | BB | / |
| 12 | *Oryza officinalis* | *OWR* | CC | / |

Note: "/" means model species (*Oryza Sativa* L. *indica* and *Oryza Sativa* L. *japonica*) used as reference genome, or wild accessions (*Oryza punctata* and *Oryza officinalis*) get from Wang's lab.

Table S2 Primers for PCR and sequencing

| ID | | Primer pair | primer | Product (bp) |
|---|---|---|---|---|
| *RCG1* | Chimerical retrogene | 1 | CGAGATTAACATTCTCATC<br>GCTTCAGGTTACAGGTTAG | 1260 |
| | | 2 | CAAAACAGCCGGATAGATAC<br>GACATTGTCTCCCATCCGAG | 1005 |
| | Parental gene | 3 | GTCCTAGAAGAAGATGGTCG<br>CCAAGATTCAACAGACAAG | 1085 |
| | | 4 | CCAATTATCCAGCGATAGTG<br>GACATTGTCTCCCATCCAAG | 1196 |
| *RCG2* | Chimerical retrogene | 1 | CTGGAAGGATGCATGGAATGG<br>CTCAGTTCCTGTAGGGCCTG | 1094 |
| | Parental gene | 2 | GAACTCCAGTTTAAAGGTTCTG<br>CACAGCTTCGAATTATCAACTC | 1912 |
| | | 3 | CAGGCAGGTGAGGTTTCCTGG<br>CTGAGTTTCTGTTCCTGATGG | 1332 |
| *RCG3* | Chimerical retrogene | 1 | GCAGCGGTACATATTGATGG<br>ACGCTGTAGACTCCATTGGG | 1377 |
| | Parental gene | 2 | TCATCCAACATGGGAAGGAG<br>ACCTTGACCTCTCCACAAGC | 1166 |
| *RCG4* | Chimerical retrogene | 1 | TGCCATCCTCACTACGAAGAC<br>TTATGTGGATCGTCTTCCGC | 1155 |
| | Parental gene | 2 | ACATGACGACTGCTTGATCG<br>TCTCTCGTCATCAAGTGAGGG | 1181 |
| *RCG5* | Chimerical retrogene | 1 | ATTCAAGGAAAGCTGGGTTG<br>GCGAAGAGACGAAGTTACCTG | 1170 |
| | Parental gene | 2 | TGGGAGATCTGGCAAAGATG<br>GCACTGTCATCGGATCATTC | 1165 |
| *RCG6* | Chimerical retrogene | 1 | GGCACTTCACCGAAGGGTAG<br>GGATCCGTTATCTCCTCTTCC | 1087 |
| | Parental gene | 2 | ATAGCGAGCAGGTCGGTTC | 1257 |

| | | | | |
|---|---|---|---|---|
| | | | ATAACAGCAGGGGCAAAGC | |
| *RCG7* | Chimerical retrogene | 1 | GGTTGAAAGGAAGAACCGC | 966 |
| | | | TAGCAACTGGTGCAAGGTTTC | |
| | Parental gene | 2 | TTCGTCTCCATGTGTTCCTC | 881 |
| | | | ACAGCATTGATCCACGATTC | |

Table S3 Chimerical retrogene and parental gene blat against IR8 genome

The sequence of chimerical retrogene and corresponding parental gene were blat searched against indica rice genome IR8, which was sequenced with Pacbio. Round brakets indicated the output of blat; angle brakets means when blat out were too long the sequences range were narrowed down by gene-specific primer.

| | Gene ID in Plant cell paper | Chimerical gene in 9311 | Chimerical retrogene in IR8 | Parental gene in 9311 | Parental gene in IR8 |
|---|---|---|---|---|---|
| *RCG1* | Chr03_4107 | Chr03:27608262-27613159 (4897) | Chr03: 26049124.. 26054020 (4896) | Chr02:14775401-14780342 (4941) | Chr02: 14335789.. 14340666 (4877) |
| *RCG2* | Chr04_4524 | Chr04:30664128-30669070 (4942) | Chr04: 31889774..31894673 (4899) | Chr04:30673408-30679130 (5722) | Chr04: 31713113..31894438 (543826) |
| *RCG3* | Chr12_904 | Chr12:6133222-6139034 (5813) | Chr12: 5723488..5727995 <4507> | Chr11:19034040-19040398 (6359) | Chr12: 5801686..5807082 (5396) |
| *RCG4* | Chr10_2602 | Chr10:17747410-17752061 (4652) | Chr10: 20792601..20812983 <3709> | Chr09:4376856-4381567 (4711) | Chr09: 5374301..5378751 (4450) |
| *RCG5* | Chr01_5436 | Chr01:36521615-36526443 (4828) | Chr01:34682823..34687645 (4822) | Chr05:18541573-18546535 (4962) | Chr05:18326109..18330814 (4705) |
| *RCG6* | Chr02_1920 | Chr02:12785385-12789823 (4438) | Chr02:12059249..12063688 (4439) | Chr07:9892475-9896854 (4379) | Chr07:10350983..10355365 (4382) |
| *RCG7* | Chr08_3454 | Chr08:24470675-24475311 (4636) | Chr08:24715472..24720109 (4637) | Chr10:11013463-11018192 (4729) | Chr10:14428416..14432001 (3585) |

Table S4 PCR sequencing of retrogenes and parental genes

| | *RCG1* | | *RCG2* | | *RCG3* | | *RCG4* | | *RCG5* | | *RCG6* | | *RCG7* | |
|---|---|---|---|---|---|---|---|---|---|---|---|---|---|---|
| | C | P | C | P | C | P | C | P | C | P | C | P | C | P |
| *Grandi* | x | x | x | 7 | x | x | x | 15 | x | 8 | x | na | x | 5 |
| *Longi* | 1 | 5 | 5 | 6 | 3 | x | 7 | 14 | 5 | x | 1 | x | 3 | 10 |
| *Alta* | x | x | x | 3 | x | 5 | 8 | 16 | x | 7 | x | na | 1 | 6 |
| *Austra* | 7 | x | x | x | x | x | x | x | 2 | x | x | x | x | 8 |
| *Rufi* | 1 | 3 | x | x | x | x | 2 | 10 | 3 | 6 | 1 | 3 | 2 | 11 |
| *Nivara a* | 1 | 4 | x | 4 | 1 | 5 | 6 | 13 | 4 | 6 | 1 | na | 3 | 9 |
| *Nivara b* | 1 | 4 | x | 4 | x | 6 | 4 | 12 | 5 | 6 | 1 | x | 3 | 9 |
| *Glab* | na | 2 | x | 2 | 4 | na | 5 | 11 | 1 | 9 | 2 | x | 3 | na |
| *Indica* | 1 | 5 | 1 | 1 | 3 | 8 | 1&* | 9 | 5 | 6 | 1 | 4* | 4&* | 7 |
| *Japonica* | na | 6 | 1 | 1 | 2 | 9 | x | x | x | x | x | x | x | x |
| *YSD8* | na | x | x | x | x | x | x | x | x | x | x | x | x | x |
| *OWR* | x | na | x | x | x | x | x | x | x | x | x | x | x | x |
| *Total sequences numbers* | 7 | | 7 | | 9 | | 16 | | 9 | | 4 | | 9 | |

C: Means the retro-chimerical gene
P: Means the parental gene
x: Means did not get PCR result
na: Means did not get valuable sequence
*: using the *Indica* reference sequence
&: The cloned sequence did not perfect match the reference sequence of 9311.
Total sequences numbers, means the number of sequence type used for phylogeny construction, which correspond to the maximum value in C and P column for each retrogene.
The different number in the two columns represent a sequence type that unique for one or several species, which correspond to the sequence number of phylogeny in Fig.2.

Table S5. The lnL value comparison and the most probable model suggestion.

| | | | | | | | | | | | |
|---|---|---|---|---|---|---|---|---|---|---|---|
| *RCG8*_AK070283 | | lnL value | p-value | | -2007.573 | -2006.7 | -2005.8 | -2004.9 | -2003.95 | -2003.4 | -2002.9 |
| method II | ORM | -2011.5071 | | -2008.60576 | 0.151(1) | 0.146(2 | 0.13(3) | 0.113(4) | 0.098(5) | 0.11(6) | 0.119(7) |
| | Optimal_Two-RM | -2008.6058 | | -2007.57293 | | 0.182(1 | 0.166(2) | 0.144(3) | 0.124(4) | 0.141(5) | 0.152(6) |
| | | | | -2006.68178 | | | 0.179(1) | 0.163(2) | 0.141(3) | 0.164(4) | 0.178(5) |
| | ORM VS TRM | 5.802594 | 1.60E-02 | -2005.7798 | | | | 0.176(1) | 0.161(2) | 0.195(3) | 0.213(4) |
| | | | | -2004.86486 | | | | | 0.177(1) | 0.238(2) | 0.262(3) |
| | | | | -2003.95414 | | | | | | 0.305(1) | 0.338(2) |
| ((((((1123_c,185_c),bb_c),cc_c),174_c)#1,193_c | | | | -2003.42738 | | | | | | | 0.291(1) |
| method III | | | | | -2004.385 | -2003.3 | -2002.9 | -2002.5 | -2002.27 | -2002.1 | -2002.1 |
| | ORM | -2011.5071 | | -2005.33736 | 0.168(1) | 0.136(2 | 0.18(3) | 0.221(4) | 0.293(5) | 0.378(6) | 0.491(7) |
| | Optimal_Two-RM | -2005.3374 | | -2004.38497 | | 0.148(1 | 0.225(2) | 0.282(3) | 0.376(4) | 0.478(5) | 0.607(6) |
| | | | | -2003.33968 | | | 0.345(1) | 0.422(2) | 0.544(3) | 0.658(4) | 0.787(5) |
| | ORM VS TRM | 12.339388 | 4.44E-04 | -2002.89386 | | | | 0.361(1) | 0.536(2) | 0.675(3) | 0.82(4) |
| | | | | -2002.47689 | | | | | 0.521(1) | 0.705(2) | 0.873(3) |
| ((((((1123_c,185_c#1),bb_c),cc_c),174_c)#1,193 | | | | -2002.27086 | | | | | | 0.593(1) | 0.865(2) |
| the final optimal model is method III two-rati | | | | -2002.12801 | | | | | | | 0.946(1) |
| *RCG9*_AK073060 | | lnL value | p-value | | -1283.652 | -1282.8 | -1282.3 | -1281.8 | -1281.47 | -1281.2 | -1281.1 |
| method II | | | | -1288.21108 | 0.003(1 *) | 0.004(2 | 0.008(3 | 0.012(4 | 0.019(5 * | 0.029(6 | 0.048(7 * |
| | | | | -1283.65187 | | 0.185(1 | 0.251(2) | 0.297(3) | 0.358(4) | 0.425(5) | 0.532(6) |
| | ORM VS TRM | #REF! | 2.59E-03 | -1282.77483 | | | 0.314(1) | 0.38(2) | 0.455(3) | 0.53(4) | 0.648(5) |
| | | | | -1282.26803 | | | | 0.338(1) | 0.449(2) | 0.541(3) | 0.677(4) |
| | | | | -1281.80854 | | | | | 0.408(1) | 0.539(2) | 0.705(3) |
| | | | | -1281.46656 | | | | | | 0.457(1) | 0.698(2) |
| ((((NIP_p,19311_p),146_p),(153_p,152_c#1)),15 | | | | -1281.18967 | | | | | | | 0.684(1) |
| method III | | | | | -1282.268 | -1281.8 | -1281.5 | -1281.2 | -1281.11 | -1281 | -1281 |
| | ii3RM vs iiiTR | 2.929036 | 8.70E-02 | -1285.11639 | 0.017(1 *) | 0.037(2 | 0.063(3) | 0.097(4) | 0.155(5) | 0.226(6) | 0.318(7) |
| | ii4RM vs iiiTR | 4.683112 | 9.62E-02 | -1282.26803 | | 0.338(1 | 0.449(2) | 0.541(3) | 0.677(4) | 0.78(5) | 0.871(6) |
| | | | | -1281.80854 | | | 0.408(1) | 0.539(2) | 0.705(3) | 0.817(4) | 0.906(5) |
| ((((NIP_p,19311_p),146_p),(153_p,152_c#2)),15 | | | | -1281.46656 | | | | 0.457(1) | 0.698(2) | 0.832(3) | 0.928(4) |
| ((((NIP_p,19311_p),146_p),(153_p,152_c#2)),15 | | | | -1281.18967 | | | | | 0.684(1) | 0.853(2) | 0.956(3) |
| | | | | -1281.1069 | | | | | | 0.696(1) | 0.926(2) |
| the final optimal models are method III 3RM、4 | | | | -1281.03031 | | | | | | | 0.989(1) |
| *RCG10*_AK069420 | | lnL value | p-value | | -853.0997 | -852.49 | -851.99 | -851.58 | -851.551 | -851.53 | -851.53 |
| method II | ORM | -854.82105 | | -853.762855 | 0.249(1) | 0.28(2) | 0.314(3) | 0.358(4) | 0.49(5) | 0.614(6) | 0.724(7) |
| | Optimal_Two-RM | -853.76286 | | -853.099711 | | 0.27(1) | 0.329(2) | 0.385(3) | 0.542(4) | 0.678(5) | 0.79(6) |
| | | | | -852.490393 | | | 0.316(1) | 0.401(2) | 0.598(3) | 0.75(4) | 0.859(5) |
| | ORM VS TRM | 2.116382 | 1.46E-01 | -851.987978 | | | | 0.365(1) | 0.646(2) | 0.821(3) | 0.921(4) |
| | | | | -851.576865 | | | | | 0.819(1) | 0.953(2) | 0.992(3) |
| | | | | -851.550692 | | | | | | 0.834(1) | 0.976(2) |
| | | | | -851.528787 | | | | | | | 0.948(1) |
| method III | ORM | -854.82105 | | | -851.9883 | -851.58 | -851.55 | -851.53 | -851.527 | -851.53 | -851.53 |
| | Optimal_Two-RM | -853.20621 | | -853.20621 | 0.119(1) | 0.196(2 | 0.346(3) | 0.5(4) | 0.645(5) | 0.763(6) | 0.85(7) |
| | | | | -851.988268 | | 0.364(1 | 0.646(2) | 0.821(3) | 0.921(4) | 0.968(5) | 0.988(6) |
| | ORM VS TRM | 3.229672 | 7.23E-02 | -851.576752 | | | 0.82(1) | 0.953(2) | 0.992(3) | 0.999(4) | 1(5) |
| | ORM VS 3RM | 5.665556 | 5.89E-02 | -851.550749 | | | | 0.834(1) | 0.976(2) | 0.997(3) | 1(4) |
| | | | | -851.528734 | | | | | 0.949(1) | 0.998(2) | 1(3) |
| | | | | -851.526683 | | | | | | 0.983(1) | 1(2) |
| NONE | | | | -851.526464 | | | | | | | 0.998(1) |
| *RCG11*-AK064415 | | lnL value | p-value | | -809.0812 | -808.89 | -808.49 | -807.99 | -807.869 | -807.87 | -807.87 |
| method II | ORM | -812.39288 | | -809.447001 | 0.392(1) | 0.575(2 | 0.592(3) | 0.572(4) | 0.676(5) | 0.789(6) | 0.87(7) |
| | Optimal_Two-RM | -809.447 | | -809.081186 | | 0.541(1 | 0.556(2) | 0.535(3) | 0.658(4) | 0.788(5) | 0.877(6) |
| | | | | -808.894312 | | | 0.371(1) | 0.404(2) | 0.562(3) | 0.727(4) | 0.842(5) |
| | ORM VS TRM | 5.891756 | 1.52E-02 | -808.493406 | | | | 0.315(1) | 0.536(2) | 0.741(3) | 0.87(4) |
| | | | | -807.989177 | | | | | 0.624(1) | 0.887(2) | 0.971(3) |
| | | | | -807.869243 | | | | | | 0.994(1) | 1(2) |
| ((((19311_p,(154_p,193_p)),1123_p),185_p),(193 | | | | -807.869218 | | | | | | | 0.995(1) |
| method III | ORM | -812.39288 | | | -809.0812 | -808.71 | -808.31 | -808.03 | -808.004 | -807.93 | -807.87 |
| | Optimal_Two-RM | -809.447 | | -809.447001 | 0.392(1) | 0.48(2) | 0.516(3) | 0.587(4) | 0.718(5) | 0.804(6) | 0.87(7) |
| | | | | -809.081186 | | 0.391(1 | 0.461(2) | 0.552(3) | 0.707(4) | 0.805(5) | 0.877(6) |
| | ORM VS TRM | 5.891756 | 1.52E-02 | -808.712577 | | | 0.368(1) | 0.507(2) | 0.701(3) | 0.814(4) | 0.891(5) |
| | | | | -808.306817 | | | | 0.459(1) | 0.739(2) | 0.859(3) | 0.928(4) |
| | | | | -808.032423 | | | | | 0.811(1) | 0.9(2) | 0.955(3) |
| ((((19311_p,(154_p,193_p)),1123_p),185_p),(193 | | | | -808.003917 | | | | | | 0.696(1) | 0.874(2) |
| the final optimal model is method III two-rati | | | | -807.927404 | | | | | | | 0.733(1) |
| *RCG12*_AK067652 | | lnL value | p-value | | -562.1073 | -561.59 | -560.99 | -560.95 | -560.89 | -560.89 | -560.89 |
| method II | ORM | -567.0609 | | -562.992726 | 0.183(1) | 0.245(2 | 0.261(3) | 0.394(4) | 0.52(5) | 0.648(6) | 0.755(7) |
| | Optimal_Two-RM | -562.99273 | | -562.107272 | | 0.308(1 | 0.327(2) | 0.509(3) | 0.656(4) | 0.786(5) | 0.875(6) |
| | | | | -561.587473 | | | 0.274(1) | 0.527(2) | 0.707(3) | 0.844(4) | 0.924(5) |
| | ORM VS TRM | 8.136352 | 4.34E-03 | -560.989917 | | | | 0.772(1) | 0.905(2) | 0.977(3) | 0.995(4) |
| | | | | -560.9478 | | | | | 0.734(1) | 0.942(2) | 0.989(3) |
| | | | | -560.890201 | | | | | | 0.945(1) | 0.998(2) |
| (((((154_c,153_c),19311_c),146_c),(Chr10_231,1 | | | | -560.88786 | | | | | | | 0.995(1) |
| method III | ORM | -567.0609 | | | -560.9899 | -560.95 | -560.89 | -560.89 | -560.888 | -560.89 | -560.89 |
| | Optimal_Two-RM | -561.66458 | | -561.664583 | 0.245(1) | 0.488(2 | 0.671(3) | 0.817(4) | 0.907(5) | 0.956(6) | 0.98(7) |
| | | | | -560.989917 | | 0.771(1 | 0.905(2) | 0.977(3) | 0.995(4) | 0.999(5) | 1(6) |
| | ORM VS TRM | 10.792638 | 1.02E-03 | -560.947724 | | | 0.734(1) | 0.942(2) | 0.989(3) | 0.998(4) | 1(5) |
| | | | | -560.890201 | | | | 0.945(1) | 0.998(2) | 1(3) | 1(4) |
| | | | | -560.88786 | | | | | 0.995(1) | 1(2) | 1(3) |
| (((((154_c,153_c),19311_c#1),146_c),(Chr10_231 | | | | -560.887838 | | | | | | 1(1) | 1(2) |
| the final optimal model is method III two-rati | | | | -560.887838 | | | | | | | 1(1) |
| *RCG13*_AK070367 | | lnL value | p-value | | -1316.55 | -1316.2 | -1316.2 | -1316.2 | -1316.17 | -1316.2 | -1316.2 |
| method II | ORM | -1322.3247 | | -1317.76875 | 0.118(1) | 0.213(2 | 0.372(3) | 0.525(4) | 0.669(5) | 0.784(6) | 0.866(7) |
| | Optimal_Two-RM | -1317.7687 | | -1316.54997 | | 0.419(1 | 0.708(2) | 0.859(3) | 0.944(4) | 0.979(5) | 0.993(6) |
| | | | | -1316.22324 | | | 0.845(1) | 0.948(2) | 0.991(3) | 0.999(4) | 1(5) |
| | ORM VS TRM | 9.11185 | 2.54E-03 | -1316.20414 | | | | 0.793(1) | 0.966(2) | 0.995(3) | 0.999(4) |
| | | | | -1316.16974 | | | | | 0.986(1) | 1(2) | 1(3) |
| | | | | -1316.16959 | | | | | | 0.986(1) | 1(2) |
| ((((152_c,19311_c),nip_c),146_c)#1,(1c1_p,nip_ | | | | -1316.16944 | | | | | | | 2(1) |
| method III | ORM | -1322.3247 | | | -1316.487 | -1316.2 | -1316.2 | -1316.2 | -1316.2 | -1316.2 | -1316.2 |
| | Optimal_Two-RM | -1316.8139 | | -1316.81387 | 0.419(1) | 0.553(2 | 0.747(3) | 0.874(4) | 0.942(5) | 0.976(6) | 0.99(7) |
| | | | | -1316.48714 | | 0.465(1 | 0.752(2) | 0.903(3) | 0.966(4) | 0.989(5) | 0.997(6) |
| | ORM VS TRM | 11.021612 | 9.01E-04 | -1316.22076 | | | 0.845(1) | 0.98(2) | 0.998(3) | 1(4) | 1(5) |
| | | | | -1316.20166 | | | | 0.968(1) | 0.999(2) | 1(3) | 1(4) |
| | | | | -1316.20085 | | | | | 1(1) | 1(2) | 1(3) |
| ((((152_c,19311_c#1),nip_c),146_c),(1c1_p,nip_ | | | | -1316.20085 | | | | | | 1(1) | 1(2) |
| the final optimal model is method III two-rati | | | | -1316.20085 | | | | | | | 1(1) |

| | | | | | | | | | | | |
|---|---|---|---|---|---|---|---|---|---|---|---|
| *RCG14*_AK068903 | | lnL value | p-value | | -928.7796 | -928.78 | -928.78 | -928.78 | -928.78 | -928.78 | -928.78 |
| method II | ORM | -943.06083 | | -929.933962 | 0.129(1) | 0.315(2 | 0.511(3) | 0.679(4) | 0.805(5) | 0.889(6) | 0.941(7) |
| | Optimal_Two-RM | -929.93396 | | -928.779625 | | 0.997(1 | 1(2) | 1(3) | 1(4) | 1(5) | 1(6) |
| | | | | -928.779618 | | | 0.996(1) | 1(2) | 1(3) | 1(4) | 1(5) |
| | ORM VS TRM | 26.253726 | 2.99E-07 | -928.779605 | | | | 1(1) | 1(2) | 2(3) | 2(4) |
| | | | | -928.779605 | | | | | 1(1) | 2(2) | 2(3) |
| | | | | -928.779605 | | | | | | 2(1) | 2(2) |
| (((((154_c,nip_c),152_c),19311_cds)#1,19311_p) | | | | -928.779606 | | | | | | | 1(1) |
| method III | ORM | -943.06083 | | | -928.7796 | -928.78 | -928.78 | -928.78 | -928.78 | -928.78 | -928.78 |
| | Optimal_Two-RM | -929.93396 | | -929.933962 | 0.129(1) | 0.315(2 | 0.511(3) | 0.679(4) | 0.805(5) | 0.889(6) | 0.941(7) |
| | | | | -928.779605 | | 0.995(1 | 1(2) | 1(3) | 1(4) | 1(5) | 1(6) |
| | ORM VS TRM | 26.253726 | 2.99E-07 | -928.779605 | | | 1(1) | 1(2) | 1(3) | 2(4) | 2(5) |
| | | | | -928.779605 | | | | 1(1) | 1(2) | 2(3) | 2(4) |
| | | | | -928.779605 | | | | | 1(1) | 2(2) | 2(3) |
| (((((154_c,nip_c),152_c),19311_cds)#1,19311_p) | | | | -928.779605 | | | | | | 2(1) | 2(2) |
| the final optimal model is method III two-rati | | | | -928.779606 | | | | | | | 1(1) |
| *RCG15*_AK070946 | | lnL value | p-value | | -1004.332 | -1003.8 | -1003.7 | -1003.7 | -1003.6 | -1003.6 | -1003.6 |
| method II | ORM | -1005.54 | | -1004.92665 | 0.276(1) | 0.321(2 | 0.474(3) | 0.643(4) | 0.753(5) | 0.851(6) | 0.915(7) |
| | Optimal_Two-RM | -1004.9266 | | -1004.3321 | | 0.298(1 | 0.517(2) | 0.725(3) | 0.832(4) | 0.917(5) | 0.962(6) |
| | | | | -1003.79117 | | | 0.627(1) | 0.888(2) | 0.943(3) | 0.984(4) | 0.996(5) |
| | ORM VS TRM | 1.226774 | 2.68E-01 | -1003.67292 | | | | 0.978(1) | 0.928(2) | 0.985(3) | 0.997(4) |
| | | | | -1003.67255 | | | | | 0.7(1) | 0.928(2) | 0.985(3) |
| | | | | -1003.59831 | | | | | | 0.988(1) | 1(2) |
| | | | | -1003.5982 | | | | | | | 2(1) |
| method III | ORM | -1005.54 | | | -1003.743 | -1003.7 | -1003.7 | -1003.6 | -1003.62 | -1003.6 | -1003.6 |
| | Optimal_Two-RM | -1003.8694 | | -1003.86941 | 0.615(1) | 0.881(2 | 0.969(3) | 0.974(4) | 0.992(5) | 0.998(6) | 0.999(7) |
| | | | | -1003.74325 | | 0.99(1) | 1(2) | 0.971(3) | 0.993(4) | 0.999(5) | 1(6) |
| | ORM VS TRM | 3.341254 | 6.76E-02 | -1003.74317 | | | 0.995(1) | 0.887(2) | 0.971(3) | 0.993(4) | 0.999(5) |
| | | | | -1003.74315 | | | | 0.624(1) | 0.888(2) | 0.971(3) | 0.993(4) |
| | | | | -1003.62321 | | | | | 2(1) | 1(2) | 2(3) |
| | | | | -1003.62415 | | | | | | 0.965(1) | 0.999(2) |
| NONE | | | | -1003.62321 | | | | | | | 2(1) |
| *RCG16*_AK073961 | | lnL value | p-value | | -1255.66 | -1255.7 | -1255.7 | -1255.7 | -1255.66 | -1255.7 | |
| method II | ORM | -1259.357 | | -1256.70924 | 0.147(1) | 0.35(2) | 0.552(3) | 0.718(4) | 0.835(5) | 0.91(6) | |
| | Optimal_Two-RM | -1256.7092 | | -1255.66022 | | 1(1) | 1(2) | 2(3) | 1(4) | 1(5) | |
| | | | | -1255.66022 | | | 1(1) | 2(2) | 1(3) | 1(4) | |
| | ORM VS TRM | 5.295444 | 2.14E-02 | -1255.66022 | | | | 2(1) | 1(2) | 1(3) | |
| | | | | -1255.66023 | | | | | 0.997(1) | 1(2) | |
| | | | | -1255.66022 | | | | | | 1(1) | |
| ((((152_c,19311_cg),193_c#1),1123_c),19311_pg); | | | | | | | | | | | |
| method III | ORM | -1259.357 | | | -1255.66 | -1255.7 | -1255.7 | -1255.7 | -1255.66 | -1255.7 | |
| | Optimal_Two-RM | -1256.7092 | | -1256.70924 | 0.147(1) | 0.35(2) | 0.552(3) | 0.718(4) | 0.835(5) | 0.91(6) | |
| | | | | -1255.66022 | | 1(1) | 2(2) | 1(3) | 1(4) | 2(5) | |
| | ORM VS TRM | 5.295444 | 2.14E-02 | -1255.66022 | | | 2(1) | 1(2) | 1(3) | 2(4) | |
| | | | | -1255.66022 | | | | 0.997(1) | 1(2) | 2(3) | |
| | | | | -1255.66022 | | | | | 1(1) | 2(2) | |
| ((((152_c,19311_cg),193_c#1),1123_c),19311_pg) | | | | -1255.66022 | | | | | | 2(1) | |
| the final optimal model is method III two-rati | | | | | | | | | | | |
| *RCG17*_AK105360 | | lnL value | p-value | | -519.9366 | -519.94 | -519.94 | -519.94 | | | |
| method II | ORM | -520.73628 | | -519.936561 | 1(1) | 1(2) | 1(3) | 1(4) | | | |
| | Optimal_Two-RM | -519.93656 | | -519.936561 | | 1(1) | 1(2) | 1(3) | | | |
| | | | | -519.936561 | | | 1(1) | 1(2) | | | |
| | ORM VS TRM | 1.599442 | | -519.936561 | | | | 1(1) | | | |
| method III | ORM | -520.73628 | | | -519.9366 | -519.94 | -519.94 | -519.94 | | | |
| | Optimal_Two-RM | -519.93656 | | -519.936561 | 1(1) | 1(2) | 1(3) | 1(4) | | | |
| | | | | -519.936561 | | 1(1) | 1(2) | 1(3) | | | |
| | ORM VS TRM | 1.599442 | | -519.936561 | | | 1(1) | 1(2) | | | |
| | | | | -519.936561 | | | | 1(1) | | | |
| NONE | | | | | | | | | | | |
| *RCG18*_AK106308 | | lnL value | p-value | | -383.5465 | -382.58 | -382.01 | -381.98 | -381.902 | -381.9 | -381.9 |
| method II | ORM | -386.58557 | | -384.700173 | 0.129(1) | 0.12(2) | 0.146(3) | 0.245(4) | 0.348(5) | 0.47(6) | 0.588(7) |
| | Optimal_Two-RM | -384.70017 | | -383.546474 | | 0.164(1 | 0.215(2) | 0.371(3) | 0.511(4) | 0.656(5) | 0.772(6) |
| | | | | -382.577474 | | | 0.287(1) | 0.549(2) | 0.717(3) | 0.853(4) | 0.93(5) |
| | ORM VS TRM | 3.7708 | 5.22E-02 | -382.010972 | | | | 0.795(1) | 0.897(2) | 0.975(3) | 0.994(4) |
| | | | | -381.97719 | | | | | 0.699(1) | 0.928(2) | 0.985(3) |
| | | | | -381.90222 | | | | | | 0.987(1) | 1(2) |
| | | | | -381.902088 | | | | | | | 0.992(1) |
| method III | ORM | -386.58557 | | | -382.0012 | -381.93 | -381.9 | -381.9 | -381.902 | -381.9 | -381.9 |
| | Optimal_Two-RM | -383.7588 | | -383.7588 | 0.061(1) | 0.161(2 | 0.294(3) | 0.446(4) | 0.591(5) | 0.715(6) | 0.812(7) |
| | | | | -382.001159 | | 0.713(1 | 0.906(2) | 0.978(3) | 0.995(4) | 0.999(5) | 1(6) |
| | ORM VS TRM | 5.653546 | 1.74E-02 | -381.933275 | | | 0.804(1) | 0.97(2) | 0.996(3) | 1(4) | 1(5) |
| | | | | -381.902548 | | | | 0.987(1) | 1(2) | 1(3) | 1(4) |
| | | | | -381.902416 | | | | | 0.996(1) | 1(2) | 1(3) |
| ((154_c,((174_c,1123_c),(146_c,Chr04_2358))#1) | | | | -381.902404 | | | | | | 0.993(1) | 1(2) |
| the final optimal model is method III two-rati | | | | -381.90236 | | | | | | | 2(1) |
| *RCG19*_AK106455 | | lnL value | p-value | | -770.6134 | -770.48 | -770.46 | -770.46 | | | |
| method II | ORM | -772.20638 | | -771.303961 | 0.24(1) | 0.44(2) | 0.638(3) | 0.792(4) | | | |
| | Optimal_Two-RM | -771.30396 | | -770.613447 | | 0.609(1 | 0.855(2) | 0.957(3) | | | |
| | | | | -770.482587 | | | 0.818(1) | 0.974(2) | | | |
| | ORM VS TRM | 1.804828 | | -770.456214 | | | | 0.999(1) | | | |
| method III | ORM | -772.20638 | | | -770.8367 | -770.82 | -770.46 | -770.46 | | | |
| | Optimal_Two-RM | -771.82448 | | -771.824484 | 0.16(1) | 0.367(2 | 0.434(3) | 0.603(4) | | | |
| | | | | -770.836731 | | 0.862(1 | 0.684(2) | 0.859(3) | | | |
| | ORM VS TRM | 0.763782 | | -770.821714 | | | 0.393(1) | 0.694(2) | | | |
| | | | | -770.457003 | | | | 1(1) | | | |
| NONE | | | | | | | | | | | |

| | | | | | | | | | | | |
|---|---|---|---|---|---|---|---|---|---|---|---|
| *RCG20*_AK108373 | | lnL value | p-value | | -654.4329 | -653.58 | -652.56 | -651.73 | -651.125 | -650.74 | -648.2 |
| method II | ORM | -659.3635 | | -655.22035 | 0.21(1) | 0.195(2 | 0.15(3) | 0.137(4) | 0.146(5) | 0.176(6) | 0.051(7) |
| | Optimal_Two-RM | -655.22035 | | -654.432911 | | 0.193(1 | 0.154(2) | 0.144(3) | 0.158(4) | 0.194(5) | 0.053(6) |
| | | | | -653.584813 | | | 0.152(1) | 0.156(2) | 0.178(3) | 0.224(4) | 0.056(5) |
| | ORM VS TRM | 8.286298 | 4.00E-03 | -652.561133 | | | | 0.197(1) | 0.238(2) | 0.303(3) | 0.069(4) |
| | | | | -651.729807 | | | | | 0.271(1) | 0.372(2) | 0.07(3) |
| | | | | -651.125151 | | | | | | 0.381(1) | 0.054(2) |
| (((((((154_c,Chr02_910),153_c),146_c),193_c), | | | | -650.740947 | | | | | | | 0.024(1 |
| method III | ORM | -659.3635 | | | -651.047 | -650.68 | -650.33 | -649.5 | -647.99 | -647.84 | -647.84 |
| | Optimal_Two-RM | -652.29016 | | -652.290155 | 0.115(1) | 0.199(2 | 0.271(3) | 0.233(4) | 0.126(5) | 0.179(6) | 0.259(7) |
| | | | | -651.046964 | | 0.389(1 | 0.489(2) | 0.378(3) | 0.191(4) | 0.267(5) | 0.378(6) |
| | ORM VS TRM | 14.146688 | 1.69E-04 | -650.67531 | | | 0.407(1) | 0.31(2) | 0.147(3) | 0.225(4) | 0.339(5) |
| | | | | -650.331917 | | | | 0.198(1) | 0.096(2) | 0.173(3) | 0.288(4) |
| | | | | -649.502565 | | | | | 0.082(1) | 0.189(2) | 0.343(3) |
| (((((((154_c,Chr02_910),153_c)#1,146_c),193_c | | | | -647.990362 | | | | | | 0.58(1) | 0.858(2) |
| the final optimal model is method III two-rati | | | | -647.837142 | | | | | | | 1(1) |
| *RCG21*_AK109583 | | lnL value | p-value | | -892.9568 | -892.83 | -892.76 | -892.72 | -892.693 | -892.69 | |
| method II | ORM | -897.93707 | | -893.224573 | 0.464(1) | 0.677(2 | 0.818(3) | 0.908(4) | 0.957(5) | 0.983(6) | |
| | Optimal_Two-RM | -893.22457 | | -892.956806 | | 0.62(1) | 0.821(2) | 0.924(3) | 0.971(4) | 0.991(5) | |
| | | | | -892.833888 | | | 0.7(1) | 0.891(2) | 0.963(3) | 0.991(4) | |
| | ORM VS TRM | 9.424984 | 2.14E-03 | -892.759599 | | | | 0.773(1) | 0.935(2) | 0.987(3) | |
| | | | | -892.717968 | | | | | 0.822(1) | 0.974(2) | |
| | | | | -892.692723 | | | | | | 0.962(1) | |
| (((193_p,19311_p),174_p),(19311_cs,19311_cg)#1); | | | | | | | | | | | |
| method III | ORM | -897.93707 | | | -892.9568 | -892.83 | -892.76 | -892.72 | -892.692 | -892.69 | |
| | Optimal_Two-RM | -893.22457 | | -893.224573 | 0.464(1) | 0.677(2 | 0.82(3) | 0.908(4) | 0.957(5) | 0.983(6) | |
| | | | | -892.956806 | | 0.62(1) | 0.824(2) | 0.924(3) | 0.971(4) | 0.991(5) | |
| | ORM VS TRM | 9.424984 | 2.14E-03 | -892.833888 | | | 0.706(1) | 0.891(2) | 0.963(3) | 0.991(4) | |
| | | | | -892.762759 | | | | 0.765(1) | 0.931(2) | 0.986(3) | |
| | | | | -892.717968 | | | | | 0.819(1) | 0.974(2) | |
| (((193_p,19311_p),174_p),(19311_cs,19311_cg)#1 | | | | -892.691743 | | | | | | 0.986(1) | |
| the final optimal model is method III two-rati | | | | | | | | | | | |
| *RCG22*_AK064216 | | lnL value | p-value | | -886.1048 | -886.1 | -886.1 | -886.1 | | | |
| method II | ORM | -890.98738 | | -886.121219 | 0.856(1) | 0.984(2 | 0.998(3) | 1(4) | | | |
| | Optimal_Two-RM | -886.12122 | | -886.10475 | | 1(1) | 2(2) | 2(3) | | | |
| | | | | -886.10475 | | | 2(1) | 2(2) | | | |
| | ORM VS TRM | 9.732318 | 1.81E-03 | -886.104751 | | | | 2(1) | | | |
| (((19311_c,nip_cg),193_c),19311_p#1) | | | | | | | | | | | |
| method III | ORM | -890.98738 | | | -886.1048 | -886.1 | -886.11 | -886.11 | | | |
| | Optimal_Two-RM | -886.12122 | | -886.121219 | 0.856(1) | 0.984(2 | 0.999(3) | 1(4) | | | |
| | | | | -886.104751 | | 0.999(1 | 2(2) | 2(3) | | | |
| | ORM VS TRM | 9.732318 | 1.81E-03 | -886.10475 | | | 2(1) | 2(2) | | | |
| | | | | -886.105959 | | | | 0.966(1) | | | |
| (((19311_c,nip_cg),193_c),19311_p#1); | | | | | | | | | | | |
| the final optimal model is method III two-rati | | | | | | | | | | | |
| *RCG23*_AK069257 | | lnL value | p-value | | -189.6534 | -189.3 | -188.84 | -188.84 | | | |
| method II | ORM | -191.37024 | | -190.534619 | 0.184(1) | 0.291(2 | 0.336(3) | 0.495(4) | | | |
| | Optimal_Two-RM | -190.53462 | | -189.653353 | | 0.4(1) | 0.445(2) | 0.653(3) | | | |
| | | | | -189.298882 | | | 0.34(1) | 0.631(2) | | | |
| | ORM VS TRM | 1.671242 | | -188.842834 | | | | 0.931(1) | | | |
| method III | ORM | -191.37024 | | | -189.6534 | -189.3 | -188.84 | -188.84 | | | |
| | Optimal_Two-RM | -190.53462 | | -190.534618 | 0.184(1) | 0.291(2 | 0.335(3) | 0.495(4) | | | |
| | | | | -189.653352 | | 0.4(1) | 0.443(2) | 0.653(3) | | | |
| | ORM VS TRM | 1.671244 | | -189.298882 | | | 0.338(1) | 0.631(2) | | | |
| | | | | -188.839126 | | | | 0.992(1) | | | |
| NONE | | | | | | | | | | | |
| *RCG24*_AK069587 | | lnL value | p-value | | -1922.324 | -1921.3 | -1920.7 | -1920.5 | -1920.01 | -1920 | -1920.2 |
| method II | ORM | -1927.311 | | -1924.38604 | 0.042(1 *) | 0.046(2 | 0.059(3) | 0.099(4) | 0.119(5) | 0.186(6) | 0.296(7) |
| | Optimal_Two-RM | -1924.386 | | -1922.32391 | | 0.152(1 | 0.191(2) | 0.298(3) | 0.327(4) | 0.46(5) | 0.635(6) |
| | | | | -1921.30011 | | | 0.261(1) | 0.442(2) | 0.459(3) | 0.626(4) | 0.812(5) |
| | ORM VS TRM | 5.849888 | 1.56E-02 | -1920.66745 | | | | 0.544(1) | 0.516(2) | 0.72(3) | 0.91(4) |
| | | | | -1920.48378 | | | | | 0.328(1) | 0.615(2) | 0.889(3) |
| | | | | -1920.00541 | | | | | | 0.899(1) | 2(2) |
| (((((1123_c,185_c),174_c),146_c#1),19311_c),19 | | | | -1919.99727 | | | | | | | 2(1) |
| (((((1123_c,185_c),174_c),146_c#1),19311_c#2),19311_p); | | | | | | | | | | | |
| method III | ORM | -1927.311 | | | -1920.791 | -1920.4 | -1920 | -1920 | -1920 | -1920 | |
| | Optimal_Two-RM | -1924.386 | | -1924.38604 | 0.007(1 *) | 0.019(2 | 0.033(3 | 0.067(4) | 0.118(5) | 0.186(6) | |
| | | | | -1920.79055 | | 0.384(1 | 0.456(2) | 0.662(3) | 0.811(4) | 0.903(5) | |
| | ORM VS TRM | 5.849888 | 1.56E-02 | -1920.41161 | | | 0.367(1) | 0.661(2) | 0.842(3) | 0.934(4) | |
| | | | | -1920.00541 | | | | 0.899(1) | 0.992(2) | 0.999(3) | |
| | | | | | | | | | 0.982(1) | 1(2) | |
| (((((1123_c,185_c),174_c),146_c#1),19311_c),19311_p); | | | | | | | | | | 1(1) | |
| (((((1123_c,185_c),174_c#2),146_c#1),19311_c#2),19311_p#2); | | | | | | | | | | | |
| the final optimal model is method III two-rati | | | | | | | | | | | |

Model fitting was optimized in OBSM (Zhang et al., 2011). *, significant at $p<0.05$; **, significant at $p<0.01$.
