## Supplemental figures for "Evolutionary patterns of the chimerical retrogenes in Oryza"

AK070283

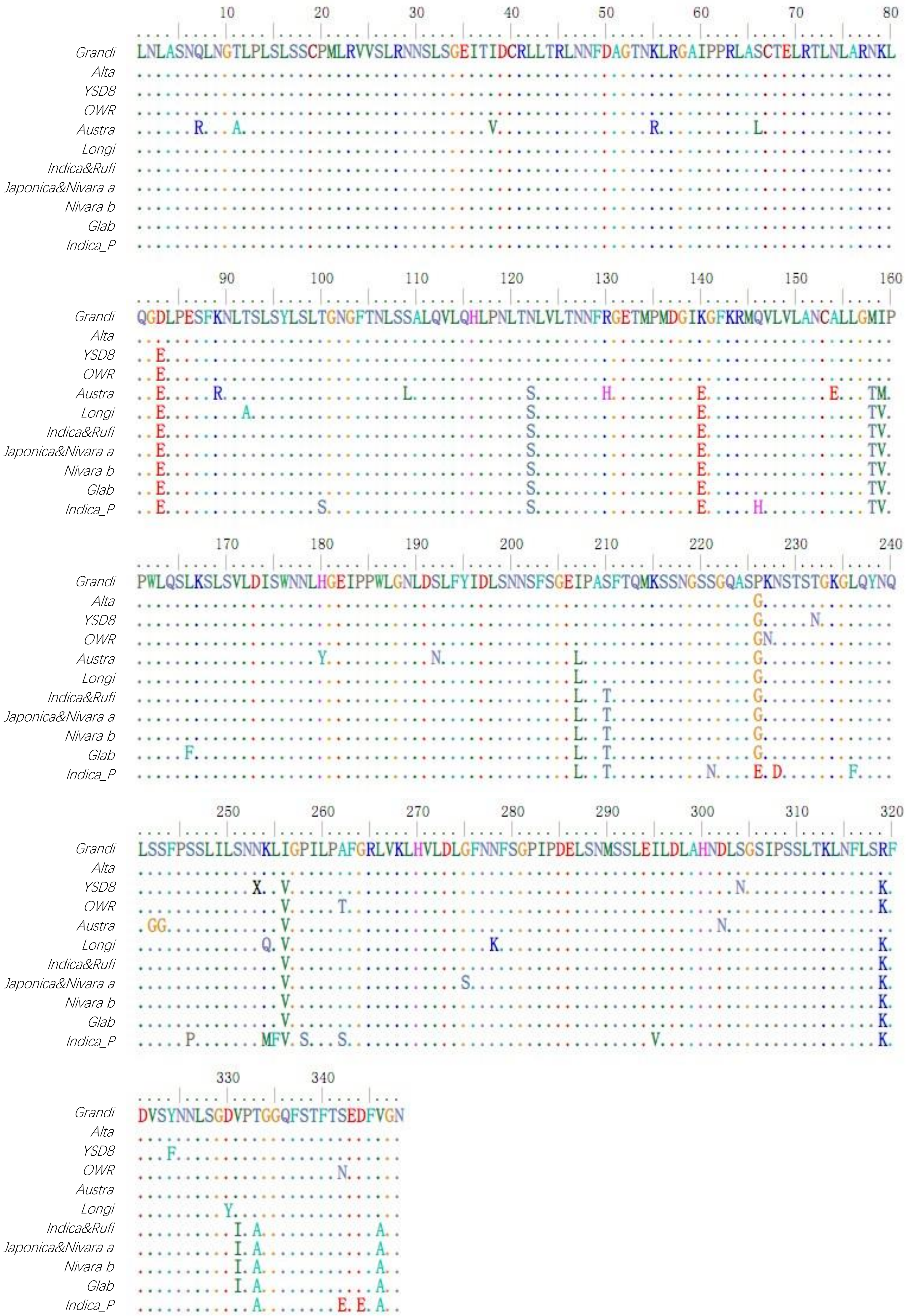
Grandi
Alta
YSD8
OWR
Austra
Longi
Indica&Rufi
Japonica&Nivara a
Nivara b
Glab
Indica_P
LNLASNQLNGTLPLSLSSCPMLRVVSLRNNSLSGEITIDCRLLTRLNNFDAGTNKLRGAIPPRLASCTELRTLNLARNKL
QGDLPESFKNLTSLSYLSLTGNGFTNLSSALQVLQHLPNLTNLVLTNNFRGETMPMDGIKGFKRMQVLVLANCALLGMIP
PWLQSLKSLSVLDISWNNLHGEIPPWLGNLDSLFYIDLSNNSFSGEIPASFTQMKSSNGSSGQASPKNSTSTGKGLQYNQ
LSSFPSSLILSNNKLIGPILPAFGRLVKLHVLDLGFNNFSGPIPDELSNMSSLEILDLAHNDLSGSIPSSLTKLNFLSRF
DVSYNNLSGDVPTGGQFSTFTSEDFVGN

AK073060

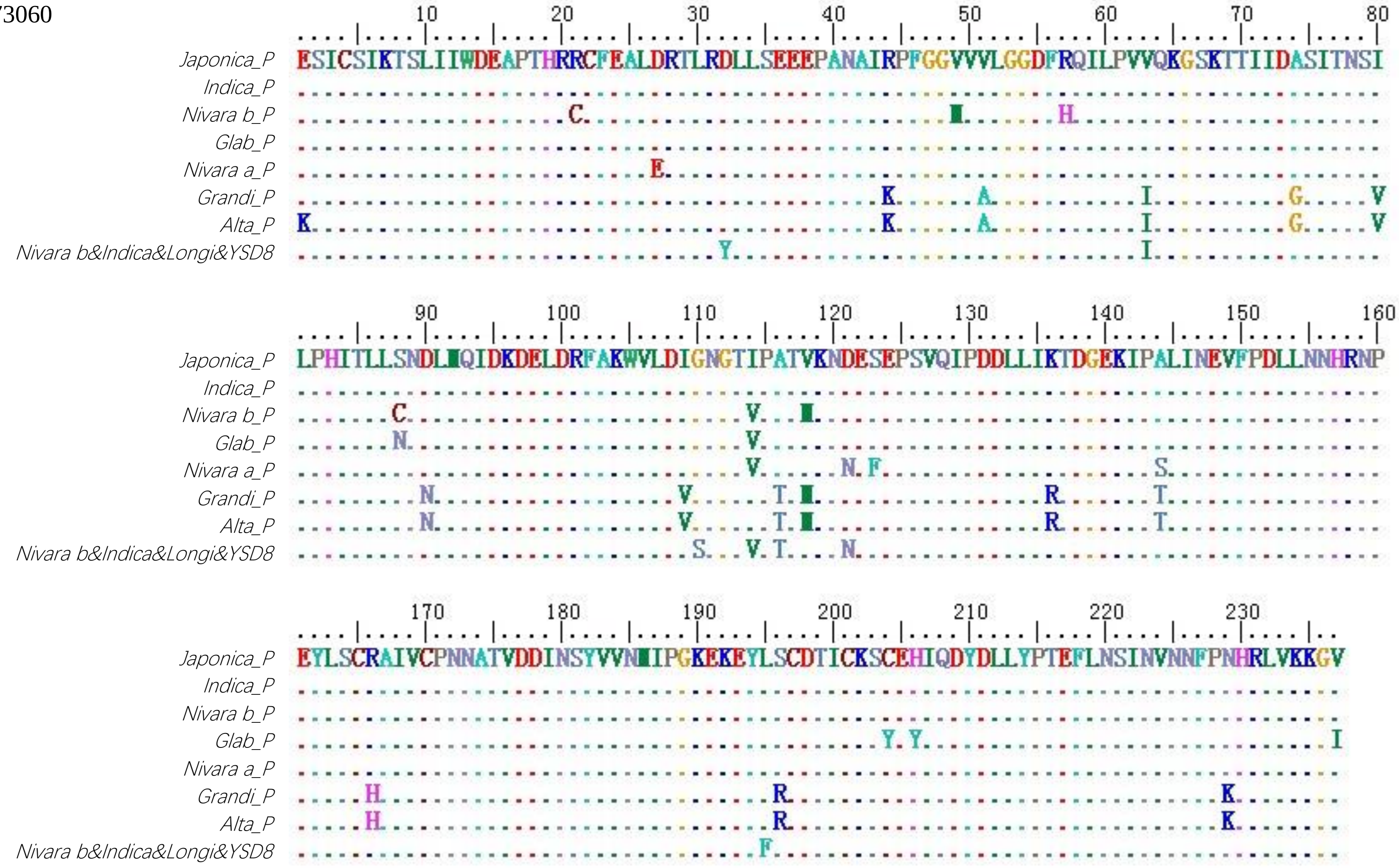


AK069420

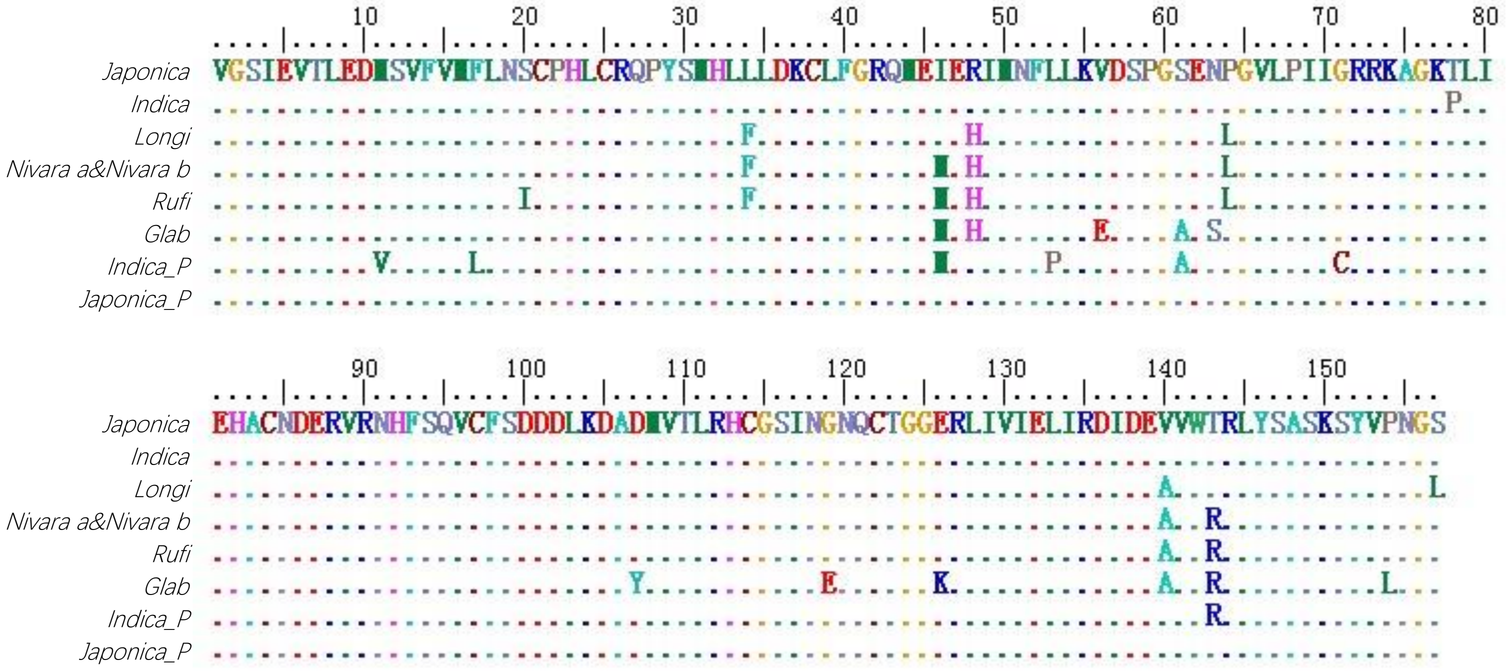


`

AK064415

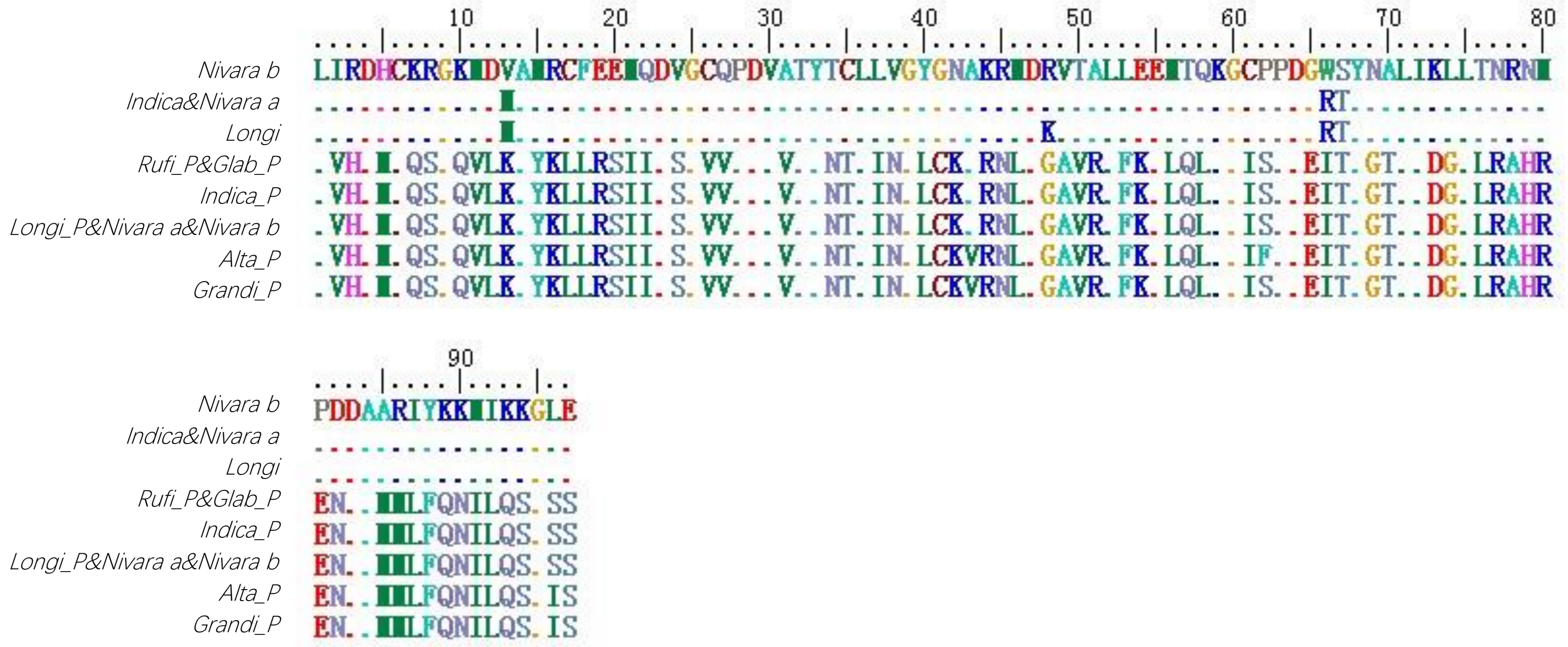


AK067652

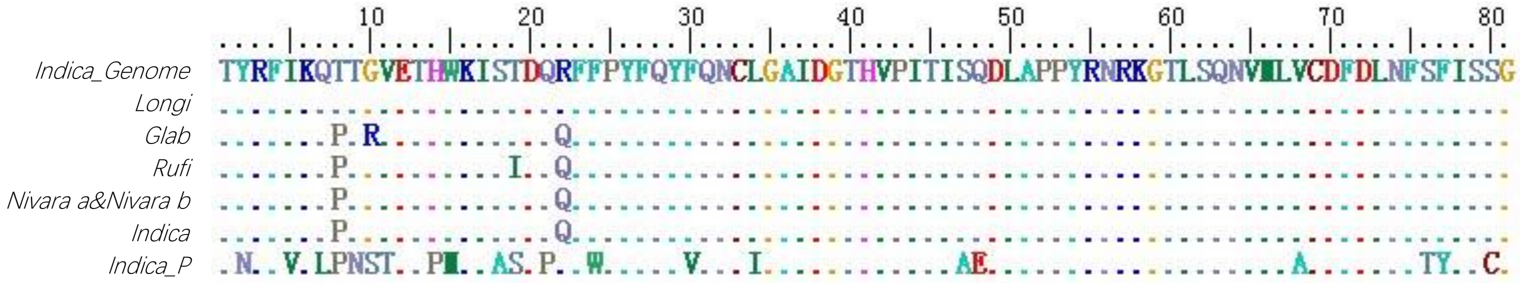


`

AK070367

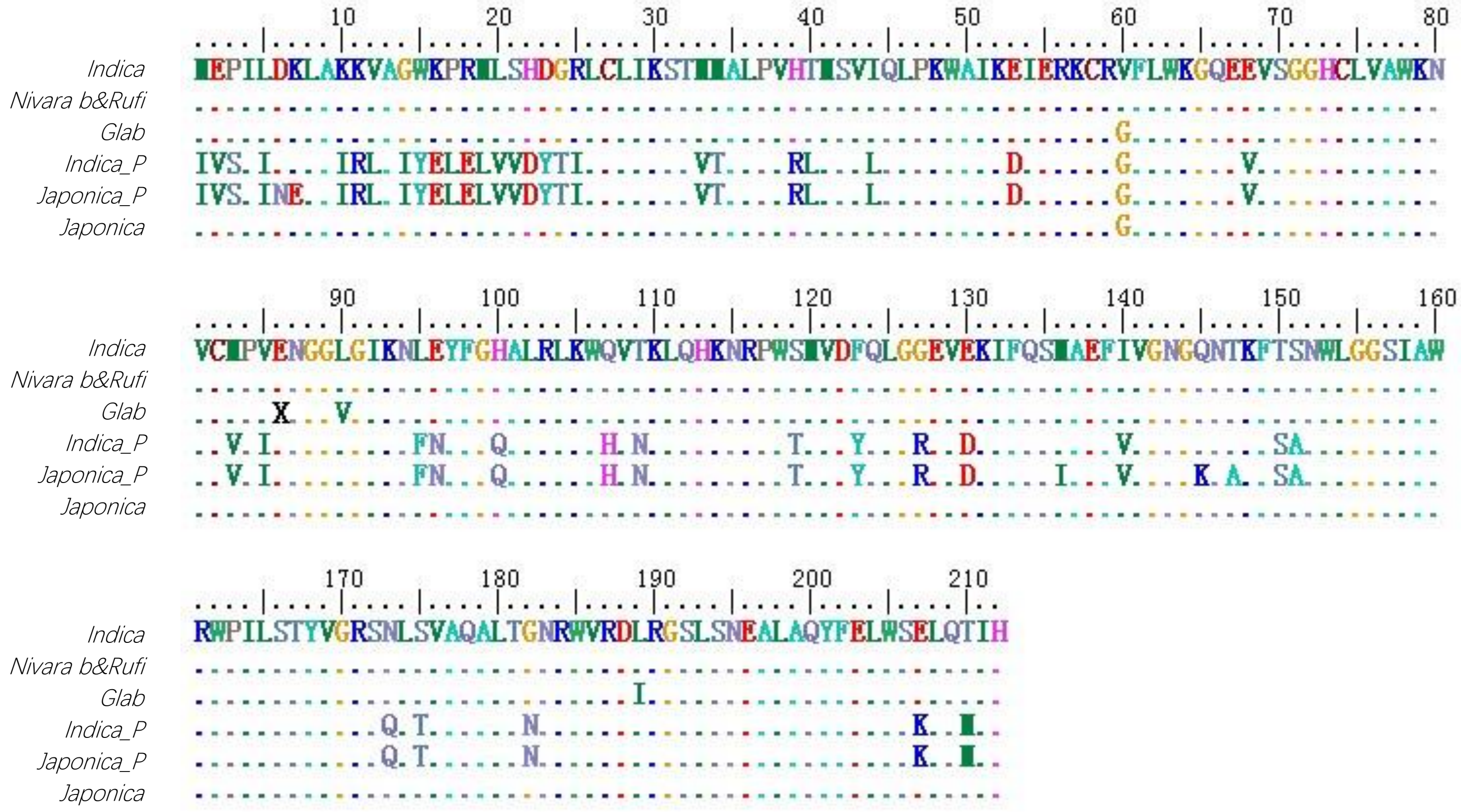


AK068903

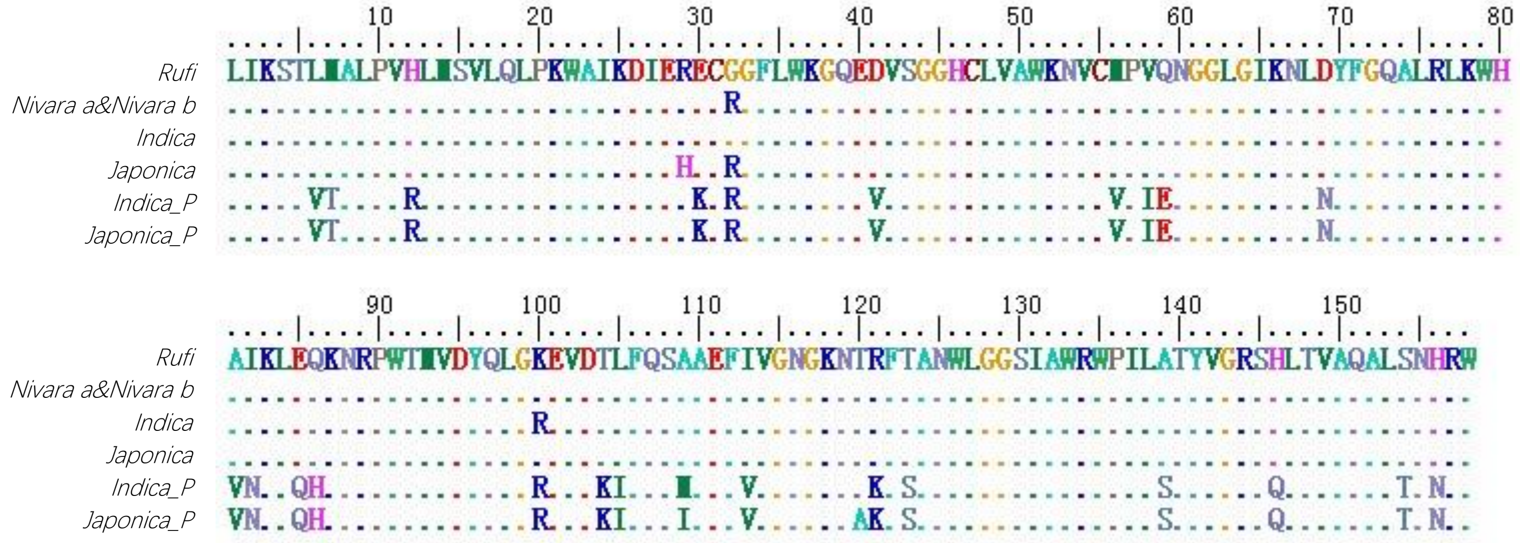


AK070946

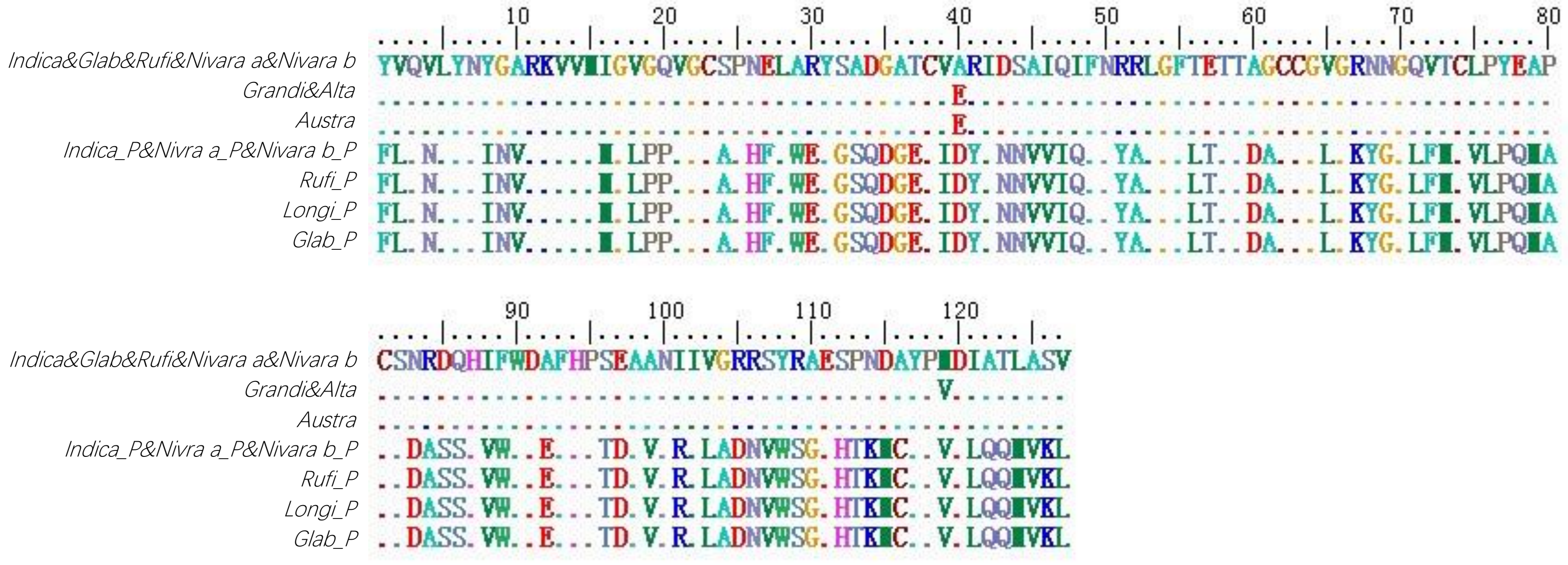

AK073961

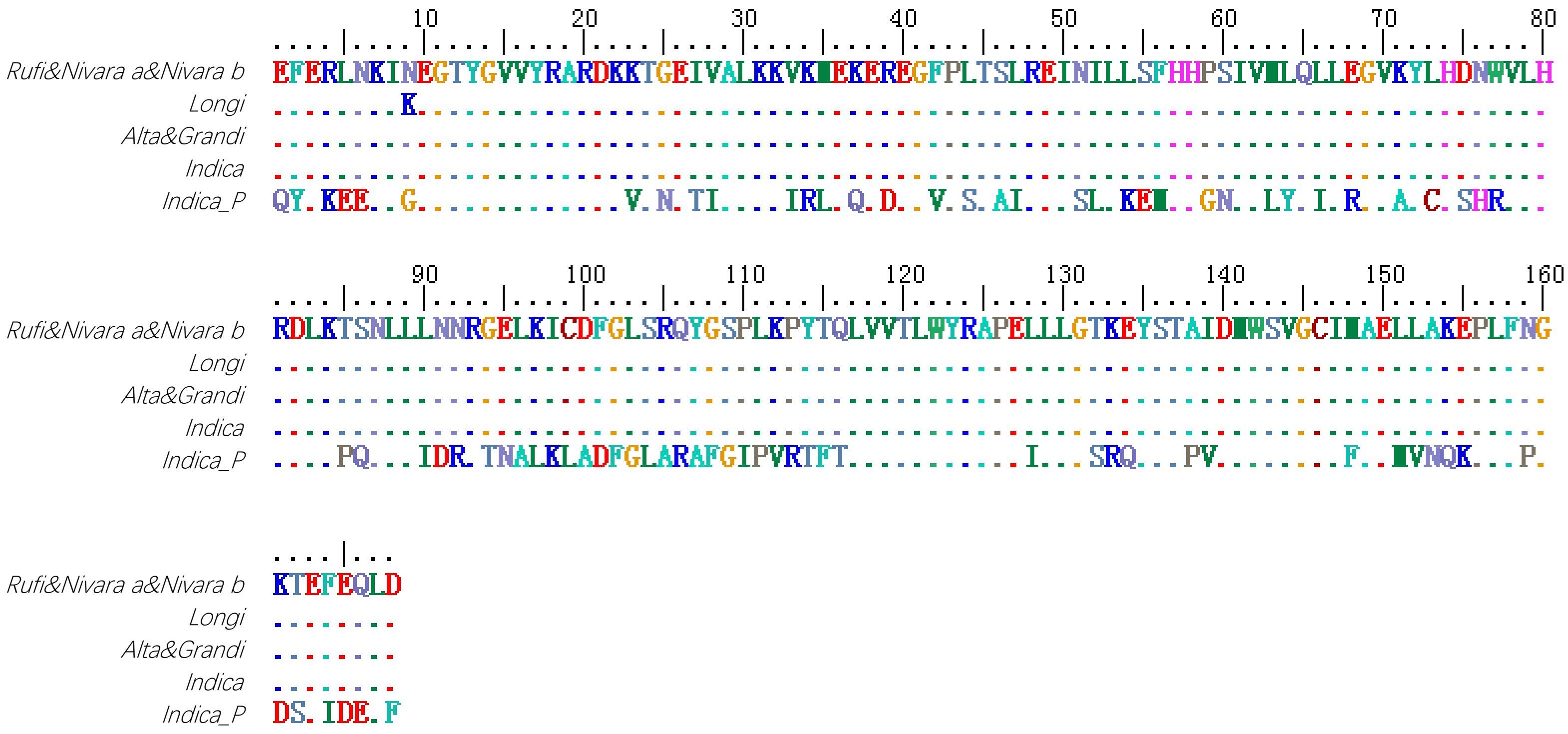


AK105360

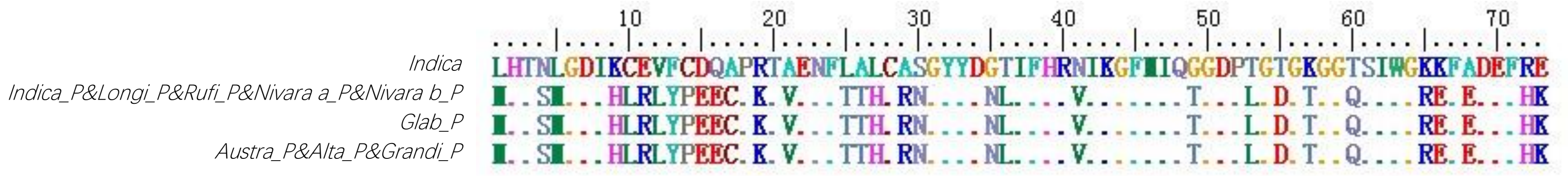


AK106308

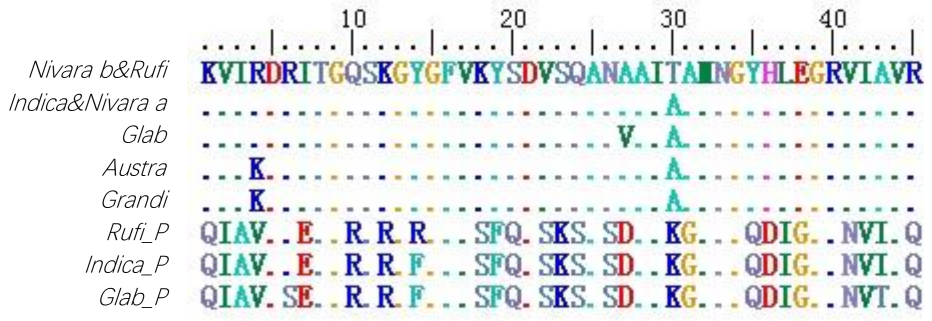


AK106455

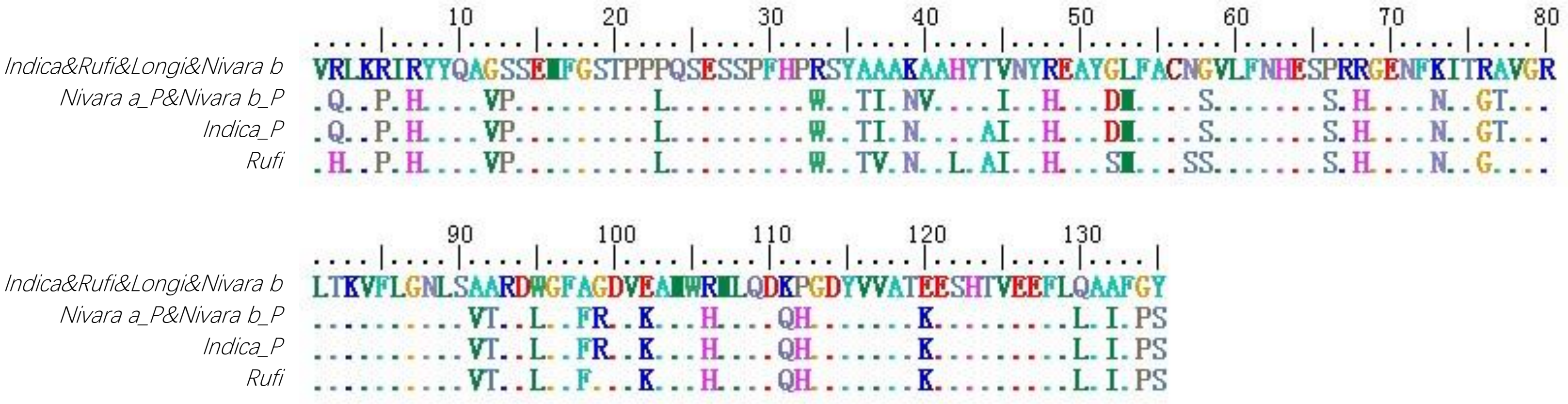


AK108373

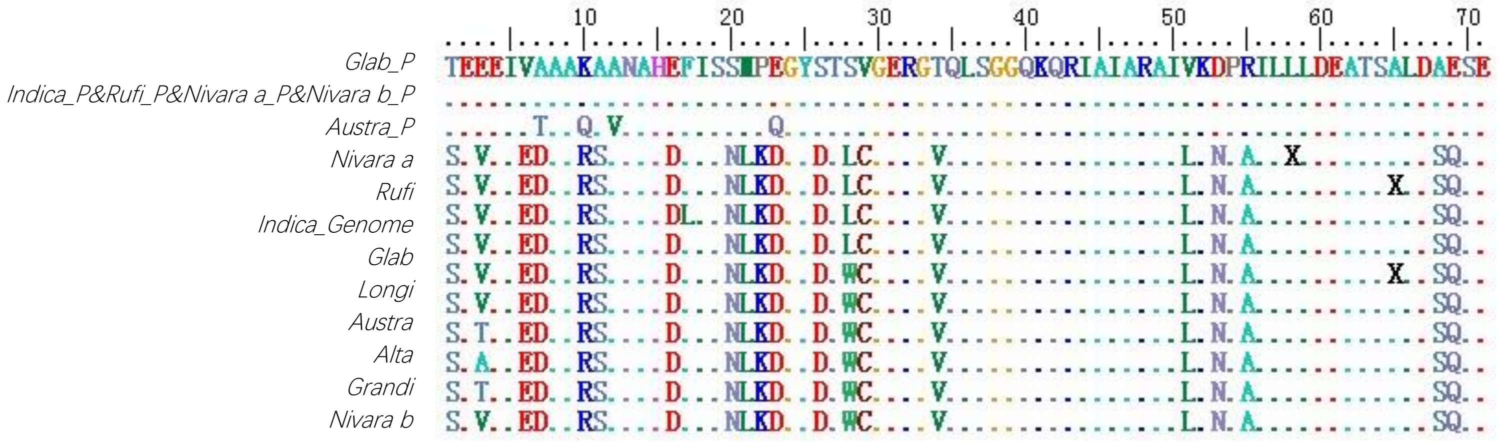

AK109583

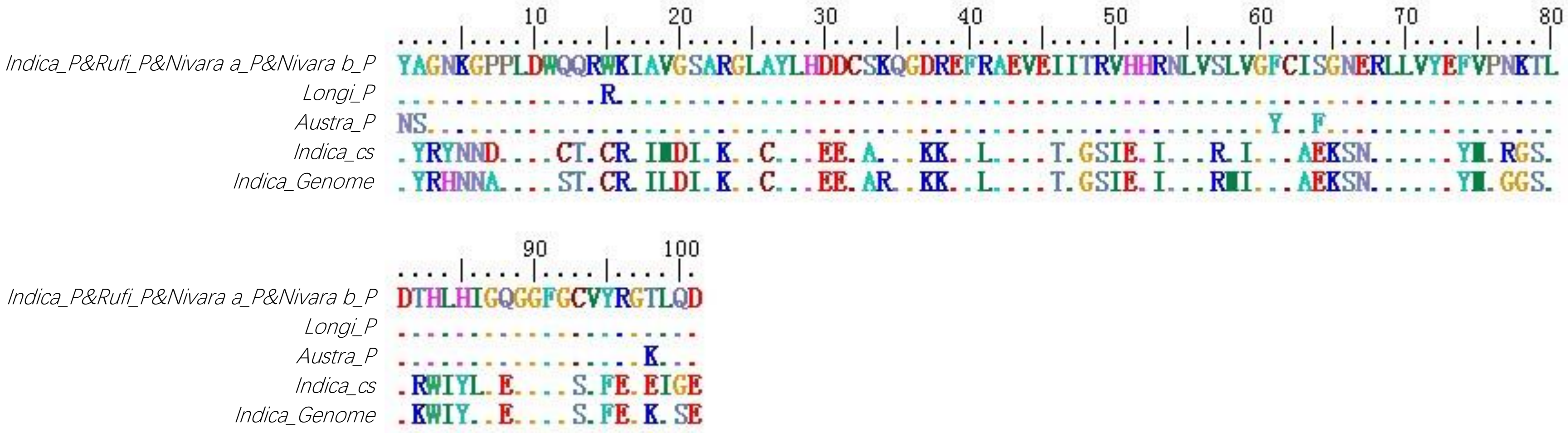


AK064216

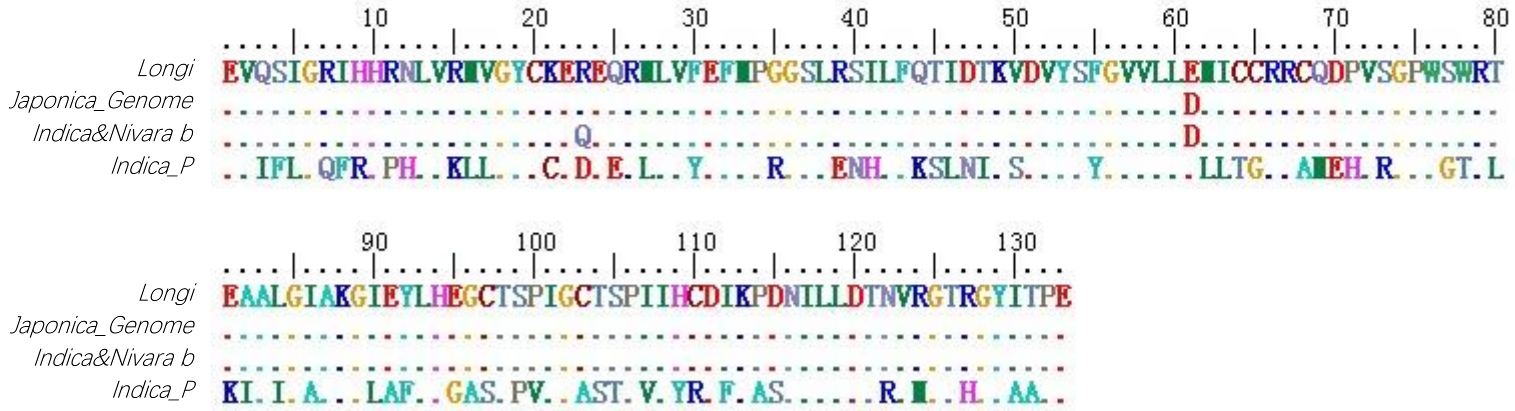


AK069257

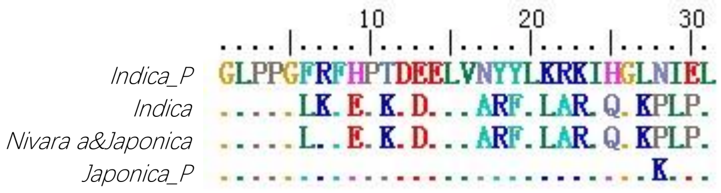


AK069587

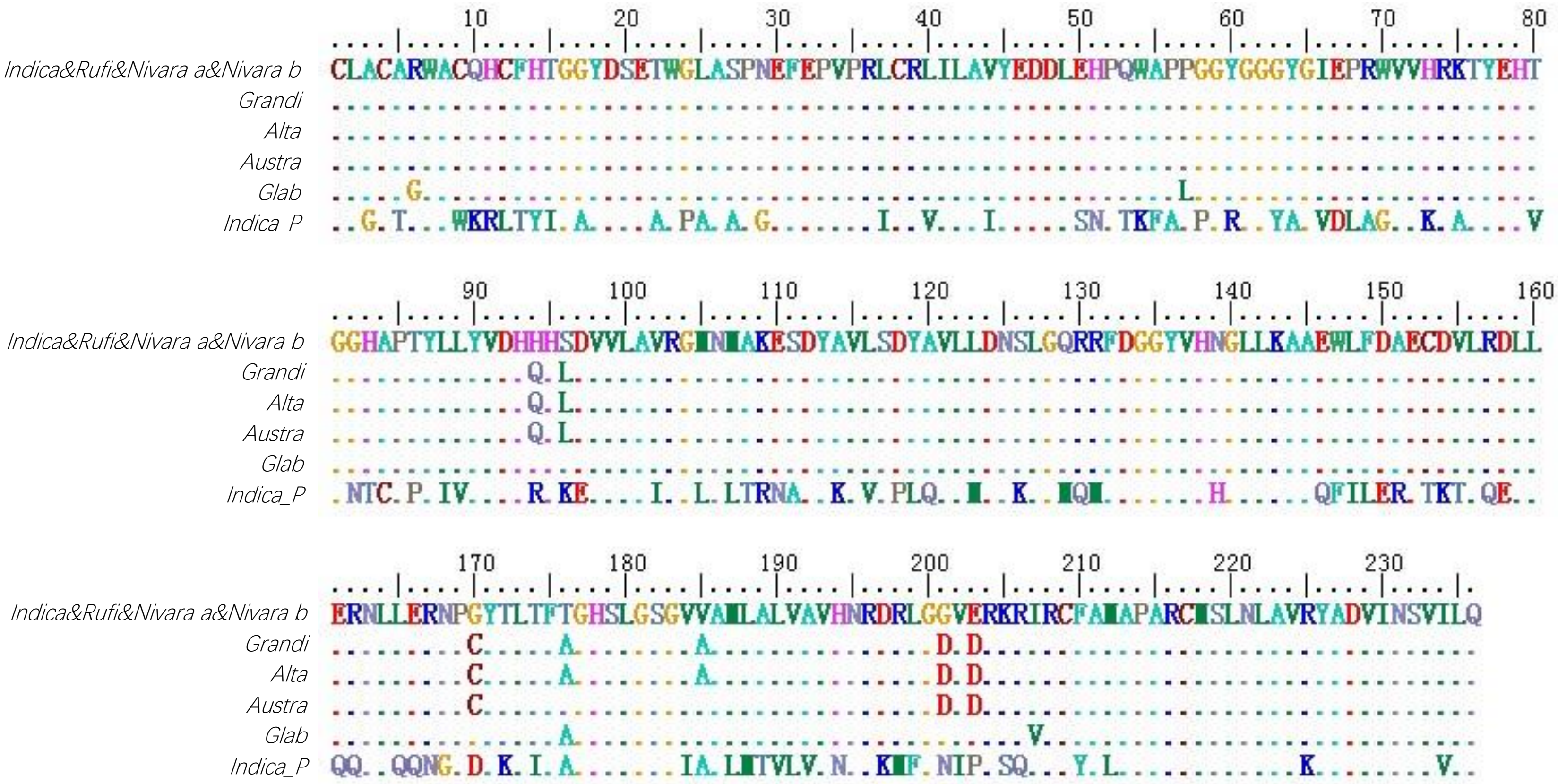


Fig. S1 The amino acid alignment of seventeen chimerical retrogene pairs.

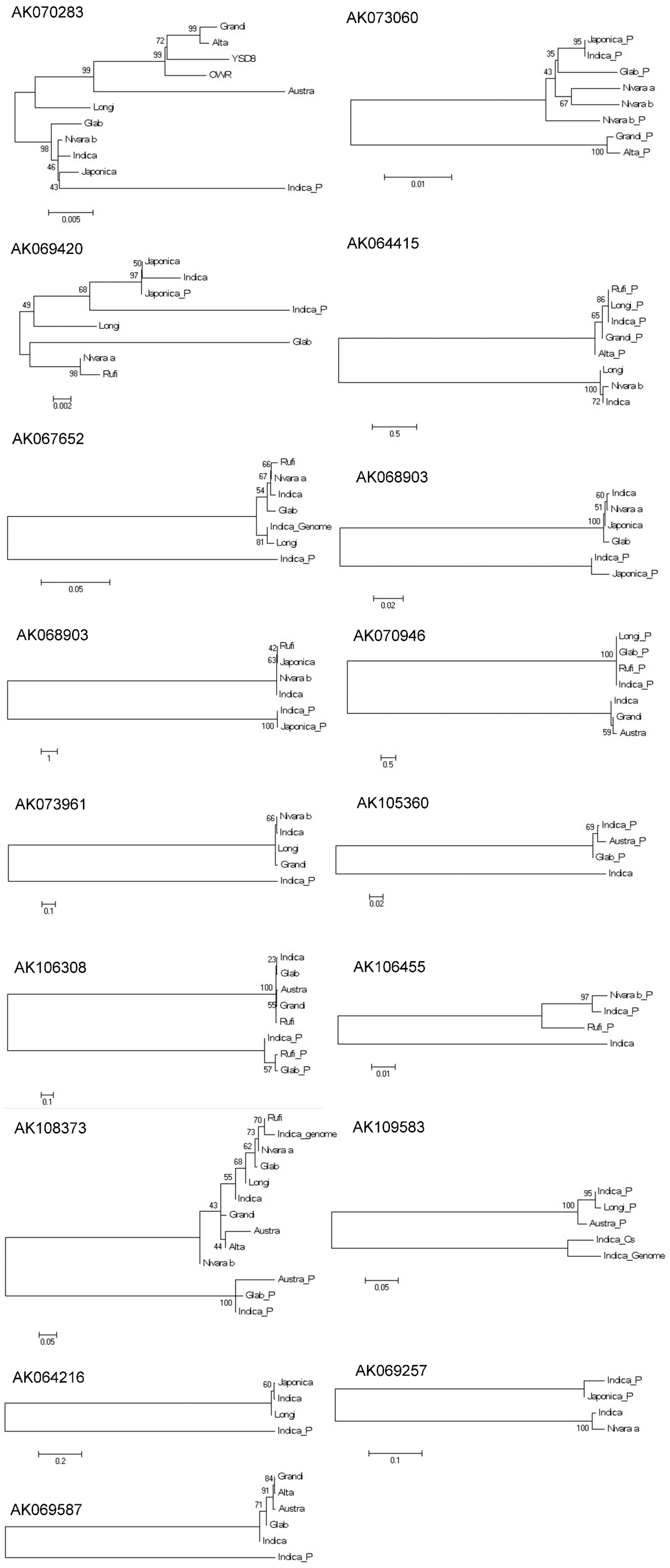


Fig. S2 The phylogeny of seventeen chimerical retrogene pairs.

AK106715

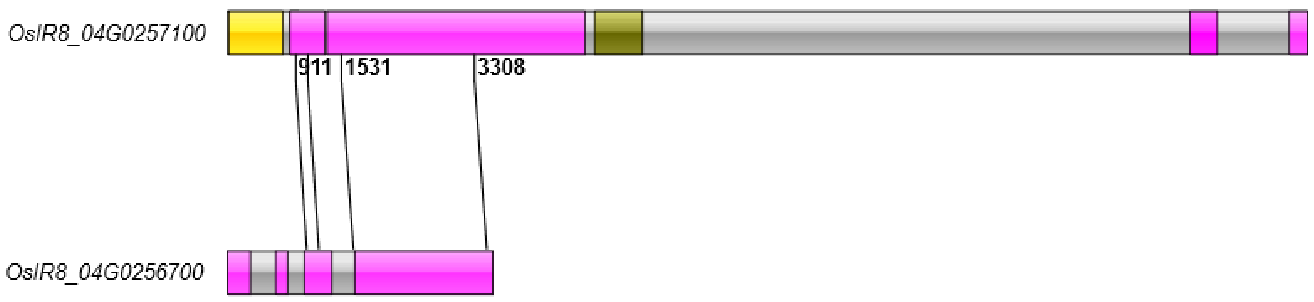


AK072107

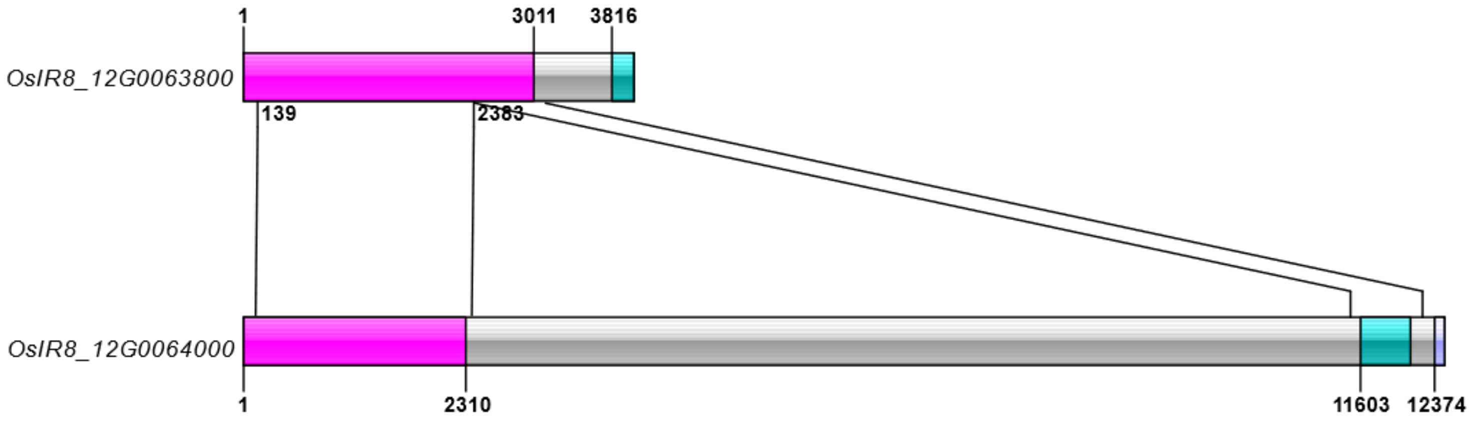


AK105722

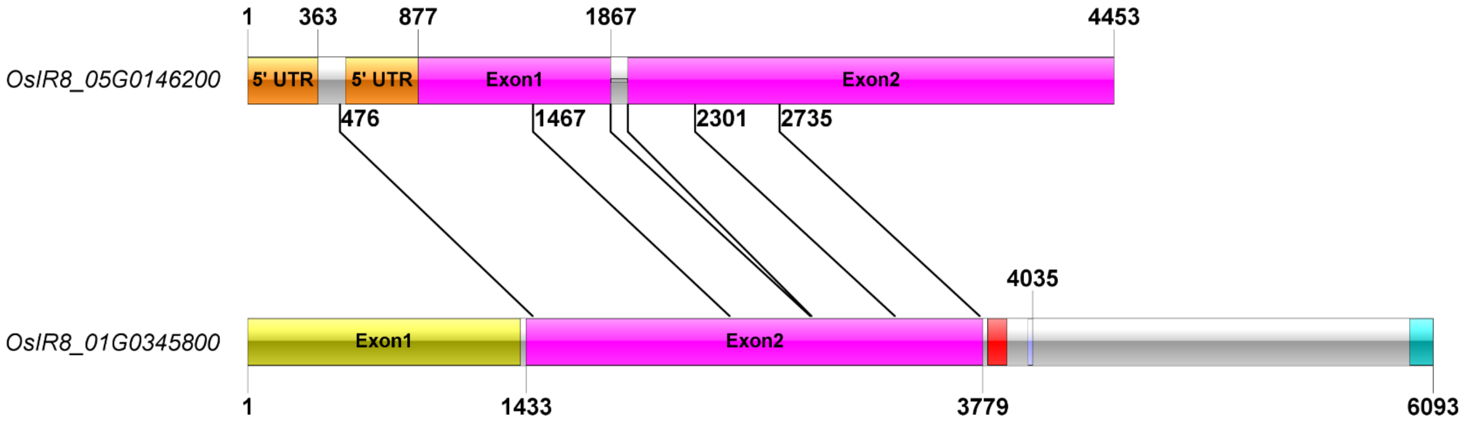


Fig. S3 Chimerical retrogene and parental gene in IR8. The sequence of chimerical retrogene and corresponding parental gene were blat searched against Indica rice genome IR8, which was sequenced by Pacbio technology. Round brackets indicated the output of blat; angle brackets mean when blat out were too long, the sequences range were narrowed down by gene-specific primer.

`
